# p53-induced apoptosis is specified by a translation program regulated by PCBP2 and DHX30

**DOI:** 10.1101/764555

**Authors:** Dario Rizzotto, Sara Zaccara, Annalisa Rossi, Matthew D. Galbraith, Zdenek Andrysik, Ahwan Pandey, Kelly D. Sullivan, Alessandro Quattrone, Joaquín M. Espinosa, Erik Dassi, Alberto Inga

## Abstract

Activation of p53 by the small molecule Nutlin can result in a combination of cell cycle arrest and apoptosis. The relative strength of these events is difficult to predict by classical gene expression analysis, leaving uncertainty as to the therapeutic benefits of Nutlin. Here, we report a new translational control mechanism shaping p53-dependent apoptosis. Using polysome profiling, we establish Nutlin-induced apoptosis to be associated with the enhanced translation of mRNAs carrying multiple copies of a newly identified 3’UTR CG-rich motif mediating p53-dependent death (CGPD-motif). We identified PCBP2 and DHX30 as CGPD-motif interactors. We found that in cells undergoing persistent cell cycle arrest in response to Nutlin, CGPD-motif mRNAs are repressed by the PCBP2-dependent binding of DHX30 to the motif. Thus, upon DHX30 depletion in these cells, the translation of CGPD-motif mRNAs is increased, and the response to Nutlin shifts towards apoptosis. Instead, DHX30 inducible overexpression in SJSA1 cells, that undergo Nutlin-induced apoptosis, leads to decreased translation of CGPD-motif mRNAs. Overall, this work establishes the role of PCBP2-DHX30 in controlling the translation of CGPD-motif mRNAs and thus provide a new mechanism to modulate the induction of p53-dependent apoptosis.

## INTRODUCTION

The tumor suppressor p53 is a tightly controlled, highly pleiotropic, stress-inducible, sequence-specific transcription factor, and is commonly inactivated in human cancer (Kruiswijk et al., 2015). Multiple regulatory circuits control p53 protein levels, localization, and activity, enabling dynamic control of its tumor suppressive functions (Kracikova et al., 2013; Sullivan et al., 2012; Vousden and Prives, 2009). An astounding amount of detail on p53-regulated transcriptional responses has been accumulated in the past three decades, yet uncertainty remains as to the critical determinants of p53 tumor-suppressive activity, particularly in solid tumors (Bieging et al., 2014).

p53 regulates an array of pathways including cell cycle arrest, DNA repair, metabolism, senescence, suppression of angiogenesis and metastasis, and modulation of innate immunity. Among these, the control of programmed cell death is often considered to be the most relevant for tumor suppression (Bieging et al., 2014). Seminal studies in mouse models, as well as evidence from the evolutionary story of the p53 pathway, have established that unrestrained p53 function can lead to massive cell death (Coffill et al., 2016; Montes de Oca Luna et al., 1995). The identification of a negative feedback loop, comprising p53 and its target and repressor MDM2 (Barak et al., 1993; Harris and Levine, 2005; Momand et al., 1992) exemplifies the evolutionary pressure to select for balanced p53 activity. It also provides a rationale to unleash p53 function as a treatment for the large fraction of cancers that retain wild-type p53 but overexpress or amplify MDM2 (Wade et al., 2013).

Several small molecules have been developed as inhibitors of the interaction between p53 and MDM2, among which Nutlin-3a (herein referred to as Nutlin) was the first and is the most extensively characterized (Khoo et al., 2014; Vassilev et al., 2004). While Nutlin-induced effects in cancer cells are indeed dependent on wild-type p53 activation, the outcome of treatment is usually a combination of cell cycle arrest, senescence, and apoptosis in relative proportions that are difficult to anticipate. This leaves uncertainty as to the potential therapeutic benefits and safety of Nutlin (Selivanova, 2014; Tovar et al., 2006). Indeed, prolonged cell cycle arrest or senescence have been associated with cancer recurrence or acquired aggressiveness (Perez-Mancera et al., 2014; Waldman et al., 1997). Consequently, many attempts have been made to untangle the pleiotropic, multifunctional p53 response, with the aim of identifying rate-limiting factors that control outcomes downstream of p53 activation. These factors could indeed be exploited as predictive or actionable markers of treatment results (Hung et al., 2011; Moumen et al., 2005; Sullivan et al., 2012). Most of those studies have focused on the regulation of p53-dependent transactivation, revealing context- and tissue-dependent cofactors that can influence the activation of pro-apoptotic p53 target genes, or shift the balance between pro- and anti-survival signals (Espinosa, 2008; Gomes and Espinosa, 2010; Gomes et al., 2006; Huarte et al., 2010; Oren, 2003; Schmitt et al., 2016). However, it is becoming evident that a conserved core of direct p53 transcriptional target genes exists. This core is similar in cancer cells of different tissues, irrespective of their phenotypic outcome, and comprises targets associated with both cell cycle arrest and apoptosis (Allen et al., 2014; Fischer, 2017; Kracikova et al., 2013; Riley et al., 2008). In other words, when focusing solely on direct p53-dependent transcriptional responses, it is not evident how to predict the propensity of different cell types to undergo apoptosis versus cell cycle arrest in response to p53 activation.

Alternatively, additional layers of regulation may modulate the p53 outcome downstream of transcriptional activation. These layers are not explored by classical gene expression analysis or ChIP-seq studies. Only recently, our previous study (Zaccara et al., 2014) and a few others have started to reveal the important role of post-transcriptional regulatory mechanisms in the regulation of p53 downstream responses (Cho et al., 2010; Loayza-Puch et al., 2013; Marcel et al., 2013, 2015; Wang et al., 2000; Yoon et al., 2012; Zaccara et al., 2014). Both RNA-binding proteins and noncoding RNAs have been implicated in these processes. However, it remains to be elucidated to what extent post-transcriptional mechanisms may impact on the cell-type-specific outcome to p53 activation.

To investigate the mechanisms underlying cell type-specific p53-dependent responses, we performed a comprehensive analysis of the transcriptome and translatome of cancer cell lines undergoing different p53-dependent outcomes. Here, we focus on mRNAs which are differentially expressed only in the polysomal fractions that could modulate phenotypic outcomes downstream of p53 activation. Polysomal profiling is based on the fractionation of cytoplasmic lysates on sucrose gradients to distinguish mRNAs that are undergoing translation from those that are not (King and Gerber, 2016).

As a proof-of-concept, we profiled total and polysome-associated mRNAs after Nutlin treatment in two cell lines that undergo different responses, cell cycle arrest in HCT116, and cell cycle arrest followed by massive apoptosis in SJSA1. Among mRNAs changed at both the total and polysomal level, we revealed the presence of a conserved subset of genes activated by p53 irrespective of the phenotypic outcome. In contrast, we observed qualitative changes in the polysome-bound mRNA repertoire: there was essentially no overlap between mRNAs differentially represented only at the translational level among the two cell lines. With the aim of discovering *cis*-elements potentially involved in differential polysome association, we identified a new motif, herein called CGPD-motif, enriched within the 3’UTR of mRNAs translationally enhanced in SJSA1 cells only. Immunoprecipitation and mass spectrometry studies indicate that in HCT116 cells the RNA helicase DHX30 along with the RNA binding protein PCBP2 bind mRNAs carrying the identified motif. PCBP2 and, to a larger extent, DHX30 knockdown in HCT116 cells lead to enhanced association of CGPD-motif mRNAs to the polysomes and to an increase in Nutlin-induced apoptosis. Depletion of DHX30 in U2OS, an osteosarcoma cell line that responds to Nutlin by cell-cycle arrest, also led to higher Nutlin sensitivity and increased protein expression of selected CGPD-motif genes. Instead, inducible overexpression of DHX30 in SJSA1 cells led to lower translation of CGPD-motif mRNAs, based on the reporter assays, polysome association, and western blotting.

Overall, these results describe a new post-transcriptional regulatory mechanism that modulates cell fate choice downstream of p53 activation. Our work thus considerably advances our understanding of how the response to such activation is defined at levels other than transcription.

## RESULTS

### The translatomes of cells undergoing apoptosis or cell-cycle arrest upon Nutlin treatment are vastly different

Since transcriptional changes caused by Nutlin treatment are invariably multifunctional (Andrysik et al., 2017), we first asked if differences in cell outcome could be explained by translatome differences. To investigate this possibility, we chose two cell lines which are known to respond differently to treatment with Nutlin as a model system. While both undergo cell cycle arrest at early time points after treatment, only in SJSA1 cells is the arrest followed by massive apoptosis. HCT116 cells instead show minimal activation of apoptosis even after prolonged treatment (Tovar et al., 2006).

We thus analyzed total and polysome-associated mRNAs 12 hours post-Nutlin treatment. This time point precedes the onset of apoptosis in SJSA1 cells, potentially avoiding indirect effects on mRNA levels. We obtained polysomal RNA fractions by sucrose gradient centrifugation of cell extracts (**Figure 1A**). Fractions up to the 80S density were considered as sub-polysomal RNA, while more dense fractions containing two or more ribosomes were considered as polysome-associated mRNAs (**Figure S1A**). Total, sub-polysomal and polysomal RNA fractions were then polyA-selected and subjected to RNA sequencing, followed by the identification of differentially expressed genes (DEGs). To characterize DEGs according to their translational status, we defined up- and down-regulated DEG classes composed of: (1) DEGs in which the change in the polysomal fraction is coupled to a change in the total fraction (Coupled); (2) DEGs that exhibit changes only in the polysomal fraction (Translationally regulated); (3) DEGs that exhibit changes in the sub-polysomal fraction, but not in the polysomal one (Unchanged in translation) **(Figure 1B, Figure S1B) (Tables S1-S2)**.

**Figure 1.**
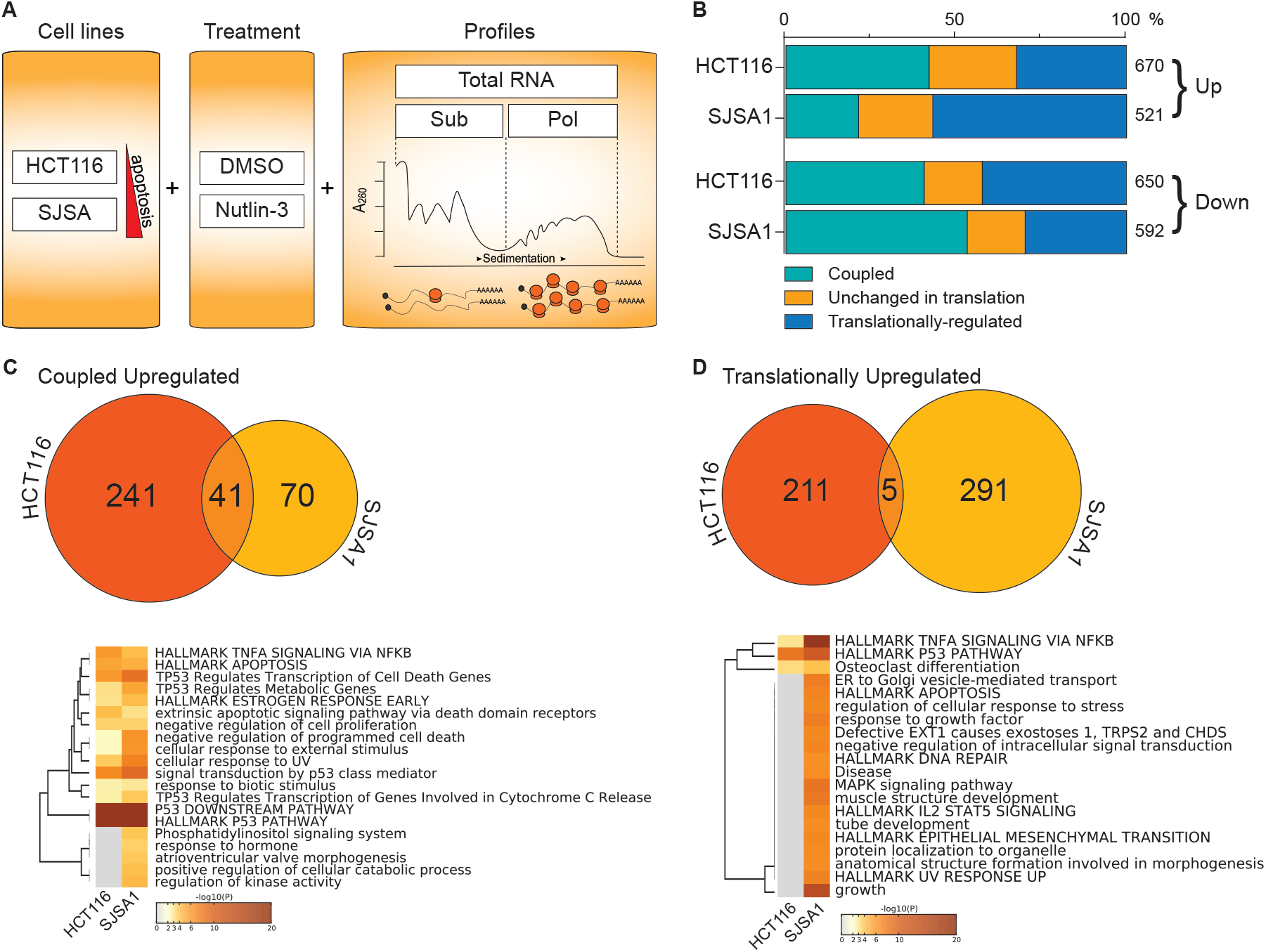
mRNA translation programs are highly cell line specific and can contribute to Nutlin-induced phenotypic outcome. **A)** Scheme of the experimental approach consisting in transcriptome (total RNA) and translatome (Sub; sub-polysomal and Pol: polysomal) analysis by RNA-seq of two p53 wild type cell lines treated with 10μM Nutlin and chosen for the distinct commitment to apoptosis (see text for details). The actual polysomal profiles after sucrose gradients fractionation are presented in Figure S1A. **B)** For each cell line, differentially expressed genes (DEGs) identified after 12-hours treatment with Nutlin were separated based on the direction of expression change (induced genes = Up; repressed genes = Down) and grouped considering the results from total RNA, cytoplasmic subpolysomal, and polysome-associated RNA. Coupled DEGs are DEGs in which the change in the polysomal fraction is coupled to a change at the total fraction (Coupled). Translationally-regulated DEGs exhibit changes only in the polysomal fraction. Unchanged in translation DEGs exhibit changes in the sub-polysomal fraction, but not in the polysomal one. The relative proportion of these three groupings and the total number of DEGs are shown. **C), D)** Top panels: Venn diagrams show the comparison of Coupled Upregulated and Translationally Upregulated DEG lists. The overlap between the two cell lines is bigger among coupled DEGs. Only 5 genes are commonly translationally upregulated. Bottom panels: Heatmaps of enriched pathways and GO terms generated using Metascape. Coupled DEGs show an enrichment of classical p53 signaling pathway terms in both cell lines. Only the translationally upregulated genes in SJSA1 cells show an enrichment of the apoptotic pathway. See also Figure S1.

Next, we compared DEGs across the two cell lines. A comprehensive analysis of the common and unique coupled DEGs has been performed separately (Andrysik et al., 2017). Briefly, in both cell lines, coupled DEGs were highly enriched for known p53-regulated targets, as confirmed by pathway and gene ontology analysis (**Figure 1C**, lower panel). The 41 commonly coupled upregulated mRNAs comprised many well-established direct p53 target genes. Importantly, as previously shown (Andrysik et al., 2017), coupled DEGs do not explain the differential outcomes following Nutlin treatment: both cell cycle arrest and apoptotic genes exhibit increases in transcription and translation across the two cell lines (**Figure 1C**, **Figure S1C**). In contrast, we noticed that, although the two cell lines show similar numbers of translationally-regulated DEGs after treatment, only 5 up-regulated and 9 down-regulated mRNAs are common to both (**Figure 1D**, **Figure S1D**).

To investigate the potential functional consequences of the distinct signatures among the translationally-regulated DEGs in the two cell lines, we carried out pathway and gene ontology enrichment analysis using Metascape (Tripathi et al., 2015). This analysis revealed the enrichment of genes associated with apoptotic signaling -Hallmark Apoptosis- (Liberzon et al., 2015) (**Figure 1D**, lower panel; **Figure S1E**, **Table S3**), only among translationally-enhanced DEGs in SJSA1 cells, potentially explaining the ability of SJSA1 cells to undergo apoptosis in response to Nutlin treatment.

Thus, while p53 deploys similar transcriptional expression programs in response to Nutlin across different cell lines, it elicits much more diverse translational programs. Consistent with our previous report (Zaccara et al., 2014), these findings highlight the importance of translationally-modulated genes in shaping phenotypic responses to p53 activation such as apoptosis.

### Cell line selective translational enhancement is mediated by a novel 3’UTR element

Given the specificity of translationally-regulated targets in each cell line, we next sought to investigate features that may determine the polysomal association of these mRNAs. A *de novo* sequence motif search (Pavesi et al., 2004) in the 5’ and 3’UTRs of DEGs identified putative cis-regulatory elements specifically enriched in the 3’UTRs of genes translationally enhanced only in SJSA1 cells (**Figure 2A**). By correlating the positional weight matrices (PWMs) of each motif in this SJSA1 gene category (**Figure S2A**, **Table S4A**), we found that three out of the eight motifs are highly similar (Pearson correlation > 0.90), corresponding to the consensus 5’-CCCC(A/C)(T/G)GGCCCT-3’, herein defined as “CGPD-motif”. By comparing the PWMs of the CGPD-motif with those identified for the other mRNA classes, we found that this motif is enriched only in the 3’UTRs of translationally enhanced mRNAs in SJSA1 cells (**Figure S2B**, **Figure S2C**, **Table S4A-B**; Pearson correlation > 0.90). Interestingly, 65% (193/296 genes) of the translationally-enhanced genes identified in Nutlin-treated SJSA1 cells harbor at least one copy of this cis-element. These 193 genes will be herein defined as “CGPD-motif genes”. Interestingly, 88.2% of CGPD-motif genes mRNAs have at least two copies in their 3’UTR. Besides a few exceptions, the expression of most CGPD-motif genes is not activated by p53 in HCT116 cells total nor polysomal RNA (**Figure S2D**).

**Figure 2.**
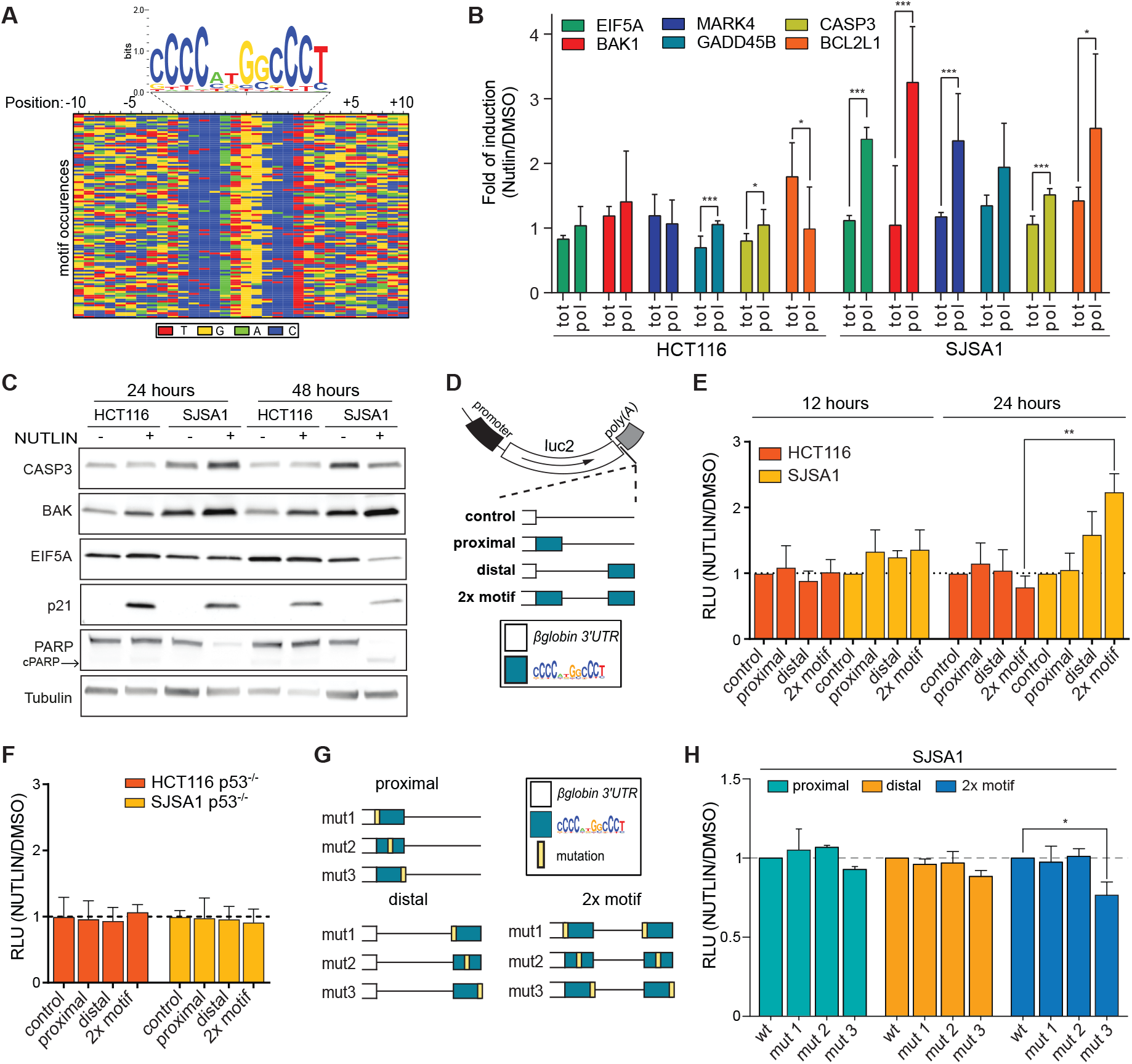
A cis-element highly enriched in 3’UTR of translationally upregulated DEGs from SJSA1 cells is sufficient to stimulate cell line specific and Nutlin-treatment dependent mRNA translation potential. **A)** Alignment for 3’UTR sequences of translationally upregulated mRNAs from SJSA1 cells centered around an enriched motif identified by Weeder (see Methods for details). The consensus sequence is presented with a logo view. **B)** RT-qPCR from total or polysomal RNA for six CGPD-motif-containing genes that were identified as translationally-enhanced in SJSA1 cells by RNA-seq. The average fold of induction and standard deviations of two biological replicates are shown. RNA was extracted from cells treated with Nutlin for 12 hours. (n=2 biological replicates each with three technical replicates, *p-value< 0.05, p-value< 0.01). **C)** Western blot analysis to detect the protein levels of the motif-containing mRNAs EIF5A, BAK and CASP3 was performed at 24 and 48 hours after Nutlin treatment. p21 was included as control of Nutlin-induced, p53-dependent coupled gene. PARP was included as control of apoptotic signaling activation. Indeed the cleaved form (indicated as c-PARP) is detectable only in SJSA1 cells upon Nutlin. Tubulin was used as loading control. CASP3 and EIF5A protein levels tend to decrease after prolonged Nutlin treatment especially in SJSA1 cells. Also, the stabilization of p21 is much reduced 48 hours after treatment. **D)** Graphical sketch of four reporter plasmids that were constructed from pGL4.13 and cloning the β -globin 3’UTR without or with the inclusion of one or two copies of the consensus CGPD-motif. The position of the motif with respect to the luciferase gene is represented. **E)** Gene reporter assays to test the potential for the identified 3’UTR motif to be sufficient in stimulating mRNA translation in the HCT116 and SJSA1 cell lines. Plasmids were transfected and the cells were then treated with Nutlin (or DMSO) for the indicated hours. Presented in the bar graphs are the average ratio of luciferase activity normalized: i) on the renilla activity; ii) for Nutlin treatment over mock; normalized to relative luciferase mRNA expression. For the 24 hour time point of panel E, separate results obtained in DMSO or Nutlin condition are presented in Figure S2F. Standard deviation of the mean are shown. **F)** p53 dependency of the impact of the CGPD-motif on reporter activity. Values were normalized as in Figure 2E. Graphical sketch **(G)** and luciferase results **(H)** in SJSA1 cells obtained with nine additional reporter plasmids developed to test the impact of three different mutations of the CGPD-motif targeting different positions of the sequence. All experiments were performed in SJSA1 cells where we observed an induction of the luciferase values when transfecting the plasmid with two copies of the motif (wt). The plasmids were transfected in SJSA1, then cells were treated with Nutlin (or DMSO) for 24 hours. Presented in the bar graphs are the average ratios of luciferase activity measured in Nutlin treated versus DMSO treated cells. Values are normalized on the RLUs detected for wt (n=3 * = p<0.05; ** = p<0.01, Student’s t-test). See also Figure S2

We validated by RT-qPCR that the translational enhancement upon Nutlin treatment occurs in SJSA1, but not HCT116 for selected CGPD-motif genes previously associated to p53-dependent cell death outcome (**Figure 2B**). Among them, BAK1, CASP3 and EIF5A were also evaluated at the protein level. BAK and CASP3 protein induction was more evident in SJSA1 cells at 24 hours of Nutlin treatment and less detectable after 48 hours. Results were less consistent for EIF5A (**Figure 2C**, **Table S15**). SJSA1 cells undergo a severe apoptotic response after 48 hours of Nutlin treatment, as confirmed by the reduction in p21 levels and the concomitant increase of the levels of cleaved PARP. We thus have identified a novel 3’UTR sequence motif, the CGPD-motif, which is enriched among translationally-enhanced mRNAs of apoptotic cells.

### The presence of the CGPD-motif is sufficient for enhancing translation in apoptosis-prone cells

To establish if the motif is sufficient for enhancing translation of CGPD-motif genes in SJSA1, one or two copies of its consensus were cloned into the β-globin 3’UTR, downstream of a firefly luciferase coding sequence, to serve as a reporter of translational efficiency. Given the absence of a preferential position for the CGPD-motif along the analyzed 3’-UTRs (**Figure S2E**), its presence at the 5’-end (proximal), the 3’-end (distal), or at both extremities (2x motif) of the 3’UTR was tested (**Figure 2D-E**). Each of the two cell lines was transfected with the reporter constructs, and luciferase activity, as well as mRNA levels, were measured. Interestingly, when placed at the distal position, a single copy of the motif led to a significant increase in reporter activity only in SJSA1 Nutlin-treated cells. Inserting two copies of the motif induced a greater effect, especially after 24 hours of treatment. Given that the reporter activity is normalized to its mRNA levels, we can rule out that changes in the reporter mRNA transcription/stability caused the induction of the 2x-motif reporter activity. On the contrary, a trend of reduction of the CGPD-motif reporter activity, even in untreated HCT116 cells is apparent, consistent with a repressive effect of the motif in this cell line (**Figure 2E**, **Figure S2F**). The enhancement of the 2x-motif reporter activity was dependent on p53 as revealed using an SJSA1 p53 knockout derivative cell line (**Figure 2F**). Moreover, when we mutated the motif at various positions, mutations at the beginning or center of the motif sequence produced luciferase levels comparable to the wild-type sequence. Since the increase was significantly reduced by modifying the last four nucleotides of the motif -here called mut3-, we conclude that the 3’ part of the sequence is essential to mediate the translational enhancement of luciferase mRNA (**Figure 2G-H**).

Thus, we have identified a novel 3’UTR-sequence motif enriched among translationally enhanced mRNAs in Nutlin-treated SJSA1 cells. This motif is sufficient to enhance the translation of a reporter mRNA in p53-expressing SJSA1 cells and confers a tendency to translational repression in HCT116 cells.

### The RNA binding proteins PCBP2 and DHX30 bind the CGPD-motif in a cell-type-dependent manner

We next set out to identify RNA-binding proteins (RBPs) capable of binding the GC-rich motif and in principle participating in the translation regulation of motif-harboring mRNAs, thus contributing to cell type-specific outcomes. As the motif is enriched among translationally enhanced mRNAs in SJSA1 but not HCT116 cells, we hypothesized that RBPs preferentially expressed or induced by Nutlin in SJSA1 cells might enhance translation of motif-containing mRNAs. Alternatively, RBPs preferentially expressed or induced in HCT116 cells could act to repress translation of CGPD-motif-containing mRNAs. We therefore aimed to identify RBPs able of recognizing the motif in a cell-line specific manner and/or having opposite expression patterns between HCT116 and SJSA1 cells. To this end, we used two different approaches. First, since the identified CGPD-motif contains a stretch of Cs, we looked for differences in the expression of PCBP proteins between SJSA1 and HCT116 cells, as they are known to make strong sequence-specific interactions with poly r(C) stretches (Leffers et al., 1995; Makeyev and Liebhaber, 2000; Matunis et al., 1992). PCBP2, but not PCBP1 or PCBP4, displays higher expression at the protein level in HCT116 compared to SJSA1 cells thus suggesting a major role of PCBP2 in the regulation of the CGPD-motif in HCT116 cells. (**Figure 3A**).

**Figure 3.**
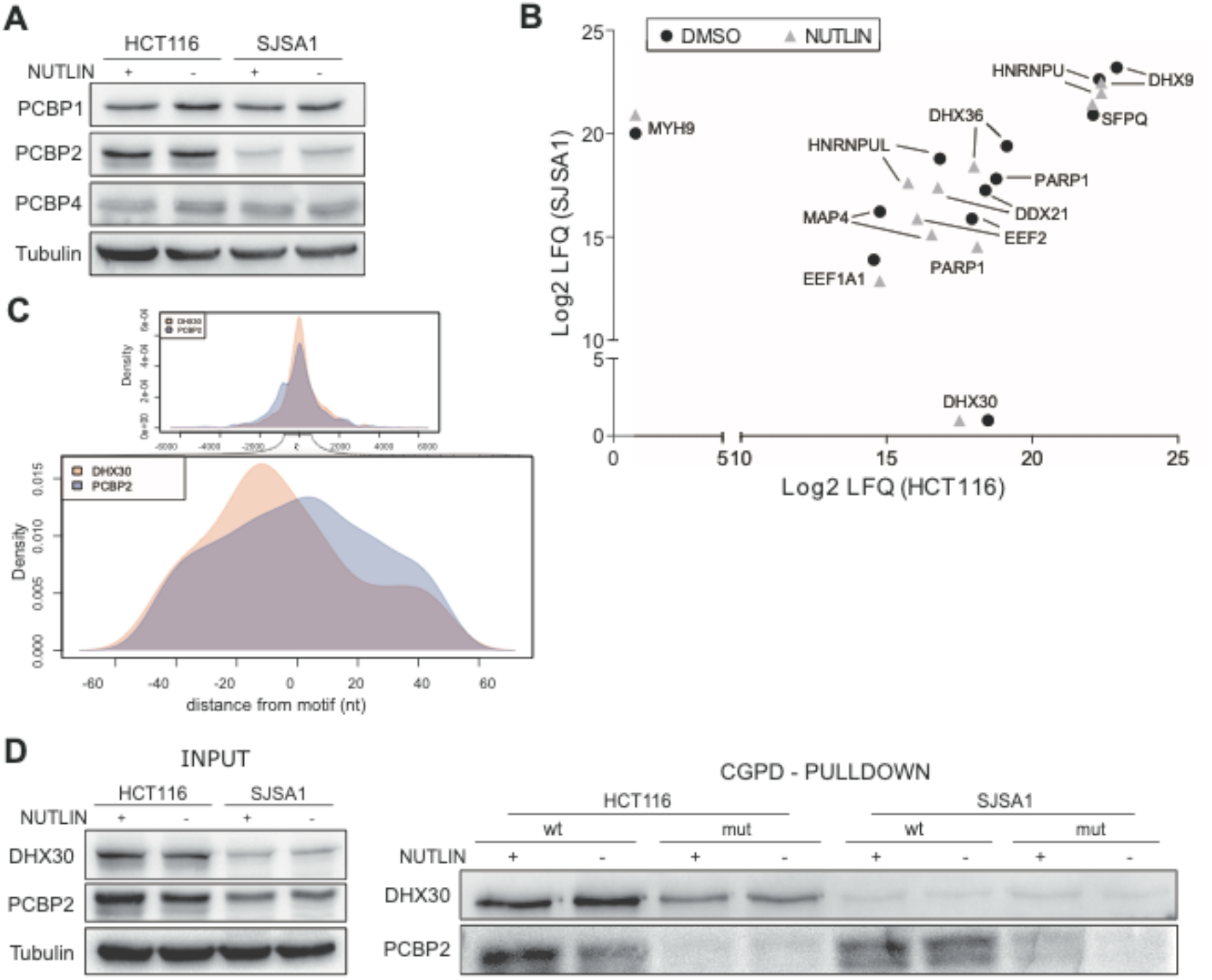
PCPB2 and DHX30 can bind the identified 3’UTR motif. **A)** Western blot of PCBP1-PCBP2-PCBP4 in HCT116 and SJSA1 cells. Given the low expression level of PCBP3 according to our RNA-seq data, we excluded PCBP3 from our analysis. Tubulin was used as loading control. **B)** Proteins from SJSA1 or HCT116 total protein extracts that were pulled-down using the consensus CGPD-motif sequence, and identified by quantitative mass spectrometry (label free quantification, LFQ). Protein extracts were prepared from DMSO or Nutlin treated cells. See **Figure S4** for additional controls of the pull-down experiment and for analysis of relative expression of DHX30 and MYH9 in HCT116 and SJSA1 cells. **C)** Positional comparison of eCLIP-derived binding sites of PCBP2 and DHX30. The graphs show the distribution of position of eCLIP sites centered on the position of the identified CGPD-rich motif, revealing a partial overlap between PCBP2 and DHX30 binding sites. **D)** Validation of PCBP2 and DHX30 interaction with the motif sequence by pull-down and immunoblot. Input is shown on the left. The pull-down was performed with the wild type CGPD-motif consensus sequence and with the mutant that showed an impact in the gene reporter assay (mut3 in **Figure 2G-H**). See also Figure S3.

Secondly, to identify RBPs in an unbiased manner, we opted for a protein pull-down approach. An RNA probe based on the motif consensus was used to capture proteins from extracts of control or Nutlin-treated HCT116 and SJSA1 cells (**Figure 3B**, **Figure S3A**). Guided by the differential bands observed by Coomassie staining, we extracted both common and specific RBPs across cell lines and treatments for identification by high-throughput quantitative mass-spectrometry analysis (**Figure 3C**; **Table S5**). Notably, we identified the RNA-helicase DHX30 as a specific hit from HCT116 extracts, while MYH9 was a specific hit from SJSA1 in at least two of the three performed experiments. Although unaffected by Nutlin treatment, DHX30 and MHY9 have corresponding patterns of differential expression at both the polysomal mRNA and protein levels (**Figure S3B**), further matching our criteria for candidate cell type-specific regulators of mRNA translation. However, the absence of an established RNA-binding domain in MYH9 sequence suggests that its binding to the CGPD-motif in SJSA1 could be indirect. We thus focus on PCBP2 and DHX30 for subsequent validation. We first re-analyzed DHX30 and PCBP2 eCLIP data in ENCODE (Van Nostrand et al., 2016). We found an enrichment for DHX30 and PCBP2 targets in our CGPD-motif-harboring mRNAs displaying enhanced translation in SJSA1 cells (Fisher p-value: p<2.2e^−16^). Conversely, the enrichment is not as significant for HCT116 translationally enhanced mRNAs (Fisher p-values: p<1.63e^−08^). Indeed, more than half of the SJSA1 translationally enhanced mRNAs (59.4%) are PCBP2 targets, compared to 25.7% of those in HCT116 cells. More importantly, we found a strong positional overlap between DHX30, PCBP2 binding sites and the CGPD-motif. As shown in **Figure 3C**, most PCBP2 binding sites in the 3’UTRs of CGPD-motif genes overlap, or are within a few nucleotides of, the motif instances, suggesting a positional interplay of PCBP2 with the CGPD-motif. Overall, these results suggest that PCBP2 and DHX30 are strong RBP candidates able to bind the CGPD-motif in vivo.

Furthermore, both DHX30 and PCBP2 are differentially expressed in HCT116 and SJSA1 -higher expression in HCT116 compared to SJSA1-further suggesting that they may contribute to the translational repression of motif-containing mRNAs in HCT116 cells (**Figure S3C**). p53 status or activation does not significantly affect MYH9, DHX30, nor PCBP2 levels (**Figure S3D**).

To validate DHX30 and PCBP2 binding to the CGPD-motif, we performed a pull-down followed by western blot analysis. As expected, binding of DHX30 to the wild-type CGPD-motif was mainly detected in HCT116 cells. In SJSA1 cells we observed only a weak immunoreactive band (**Figure 3D**). Binding of PCBP2 with the wild-type CGPD-motif was instead detected in both cell lines (**Figure 3D**). Notably, the binding of DHX30 in HCT116 and, particularly, of PCBP2 in both cell lines was sensitive to the mutation of the motif (mut3, defined in Figure 2G), suggesting that the interaction is specific.

Taken together, we found that the complex of RBPs binding the CGPD-motif is different in the two cell lines. We suggest that PCBP2 acts as a tethering factor at the CGPD-motif, since its binding is sequence-dependent but cell line-independent. Cell type-specific expression of DHX30 could act to reduce the translational efficiency of CGPD-motif mRNAs in HCT116 cells. However, considering the relative impact of sequence mutation on DHX30 binding, we cannot exclude that DHX30 can bind the motif also independently from PCBP2. We can thus suggest that the binding of these two proteins is potentially involved in the translational control of mRNAs with the CGPD-motif.

### RNA-dependent cooperation of PCBP2 and DHX30 on the CGPD-motif in HCT116 cells

The overlapping profiles of PCBP2 and DHX30 binding sites at the CGPD-motif genes 3’UTRs, along with the relative expression patterns in HCT116 and SJSA1 cells, suggest the possibility of a functional interaction between PCBP2 and DHX30 at the CGPD-motif site. To study this interaction, we took advantage of HCT116 cells given the higher levels of PCBP2 and DHX30 proteins and their stronger binding to the CG motif. PCBP2 interacts with DHX30 in an RNA-dependent manner, as demonstrated by *in vivo* co-immunoprecipitation assays (**Figure 4A**)

**Figure 4.**
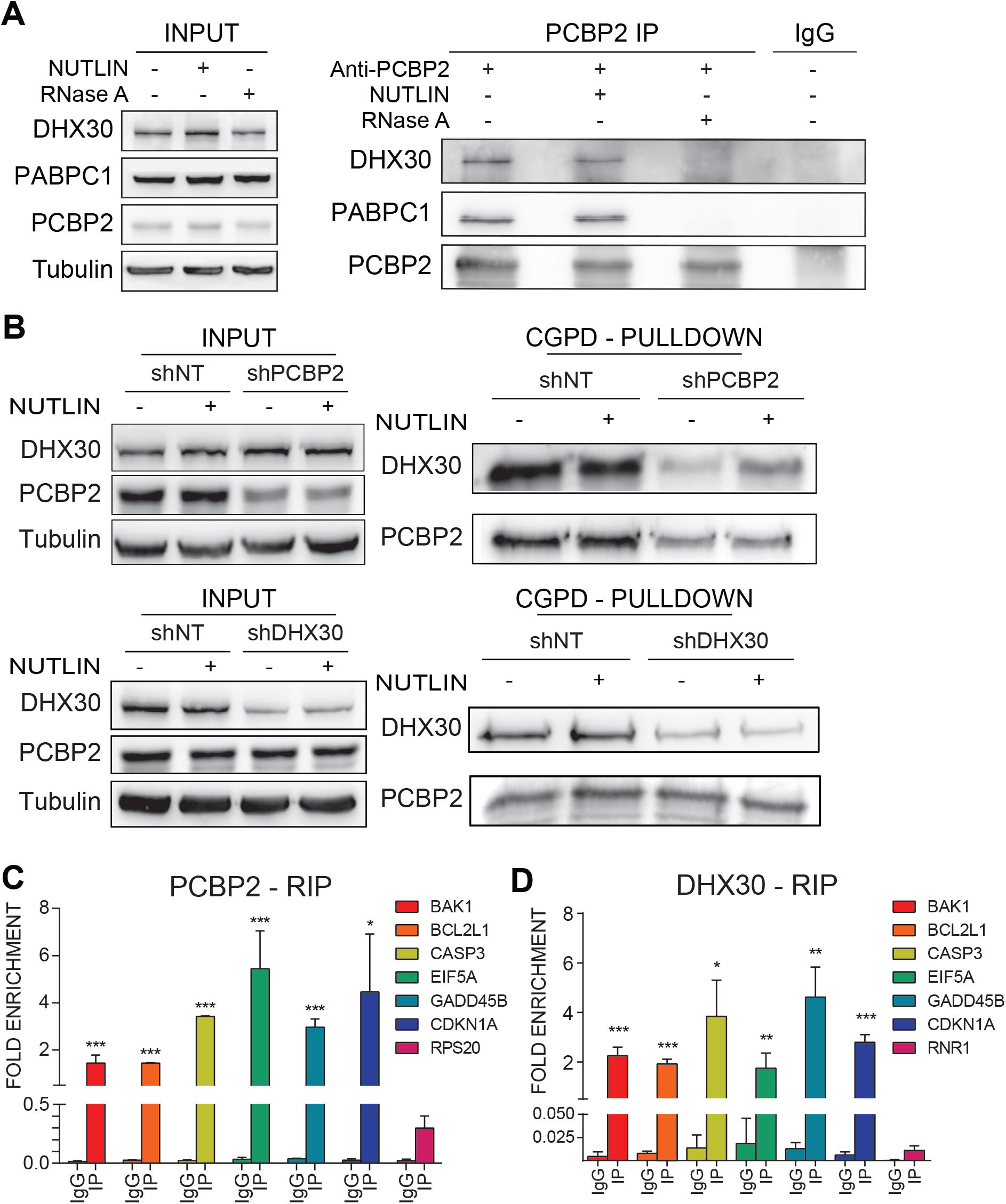
PCBP2 silencing impairs DHX30 binding to the identified 3’UTR motif. **A)** Co-immunoprecipitation of PCBP2 and DHX30 in the HCT116 parental line untreated or treated with Nutlin. RNAse A was included to test the dependency of the protein interaction on RNA. PABPC1 was tested as a reported RNA-dependent interactor of PCBP2 ^75^. **B)** The pull-down using the wild type consensus motif as bait (see Figure 3D) was repeated using HCT116 derivative clones after stable silencing of PCBP2 or DHX30. Western Blot of input proteins confirms the reduced expression only of the intended target. PCBP2 depletion impairs DHX30 binding (pull-down on top right), while DHX30 depletion has no effect on PCBP2 binding (pull-down on bottom right). RNA immuno-precipitation (RIP) assays using a PCBP2 (**C**) or a DHX30 (**D**) antibody in HCT116 cells. Data are plotted as % of input fold enrichment relative to the signal obtained for each mRNA examined in the control immunoprecipitation (IgG). Bars plot the average and the standard deviation of three qPCR replicates. (**, p<0.01, Student’s t-test). The experiments were repeated twice. As suggested by eCLIP data (see Figure 3B), PCBP2 binds motif-containing mRNAs in HCT116 cells (Wan et al., 2015). See also Figure S4.

We thus depleted PCBP2 or DHX30 and then tested their respective binding, at first *in-vitro* using the CGPD-motif RNA probe. As described in the Methods section (see also **Figure S4A**), complete knock-out of these proteins was not attainable, hence we exploited stable single-cell derived clones expressing shRNAs, obtaining comparable results on at least two different clones. When testing DHX30 binding in PCBP2-depleted cells, we observed that the DHX30 interaction with the motif was strongly reduced compared to control HCT116 cells (**Figure 4B**). In DHX30-depleted cells, DHX30 binding was nearly lost without affecting PCBP2 binding. Overall, these data suggest that DHX30 is dependent, at least in part, on PCBP2 for its binding to the CGPD-motif, while PCBP2 binding is independent from DHX30.

We then validated the interaction of PCBP2 and DHX30 on endogenous CGPD-motif mRNAs by means of RNA immunoprecipitation (RIP). As suggested by eCLIP-data, PCBP2 binds specifically to selected CGPD-motif mRNAs (**Figure 4C**). Interestingly, in DHX30 depleted cells, PCBP2 still efficiently and specifically binds CGPD-motif mRNAs, thus supporting the *in vitro* results (**Figures S4B**). As expected, significant binding to the previous CGPD-motif mRNAs was detected also when we immunoprecipitated DHX30 (**Figure 4D**). Immunoprecipitation controls for the PCBP2 and DHX30 antibodies are shown in **Figure S4C-S4D**.

Collectively, our results demonstrate an RNA-dependent cooperation between PCBP2 and DHX30 that could affect the translational efficiency of specific mRNAs.

### DHX30 and PCBP2 depletion enhances mRNA polysome association in Nutlin-treated HCT116 cells

To establish if PCBP2 or DHX30 modulate the translation of CGPD-motif-containing mRNAs, we first tested the motif-containing reporters in HCT116 cells with knockdown of either PCBP2 or DHX30 (**Figure 5A**, **Figure S5A**). While the reporter activity did not change in response to Nutlin in control cells (shNT), confirming the results of parental HCT116 cells (**Figure 2**), the depletion of both PCBP2 and DHX30 led to a significant increase in the reporter activity upon Nutlin treatment (at 12 hours for shPCBP2 cells, and at 24 hours for shDHX30 cells; data normalized to luciferase mRNA levels).

**Figure 5.**
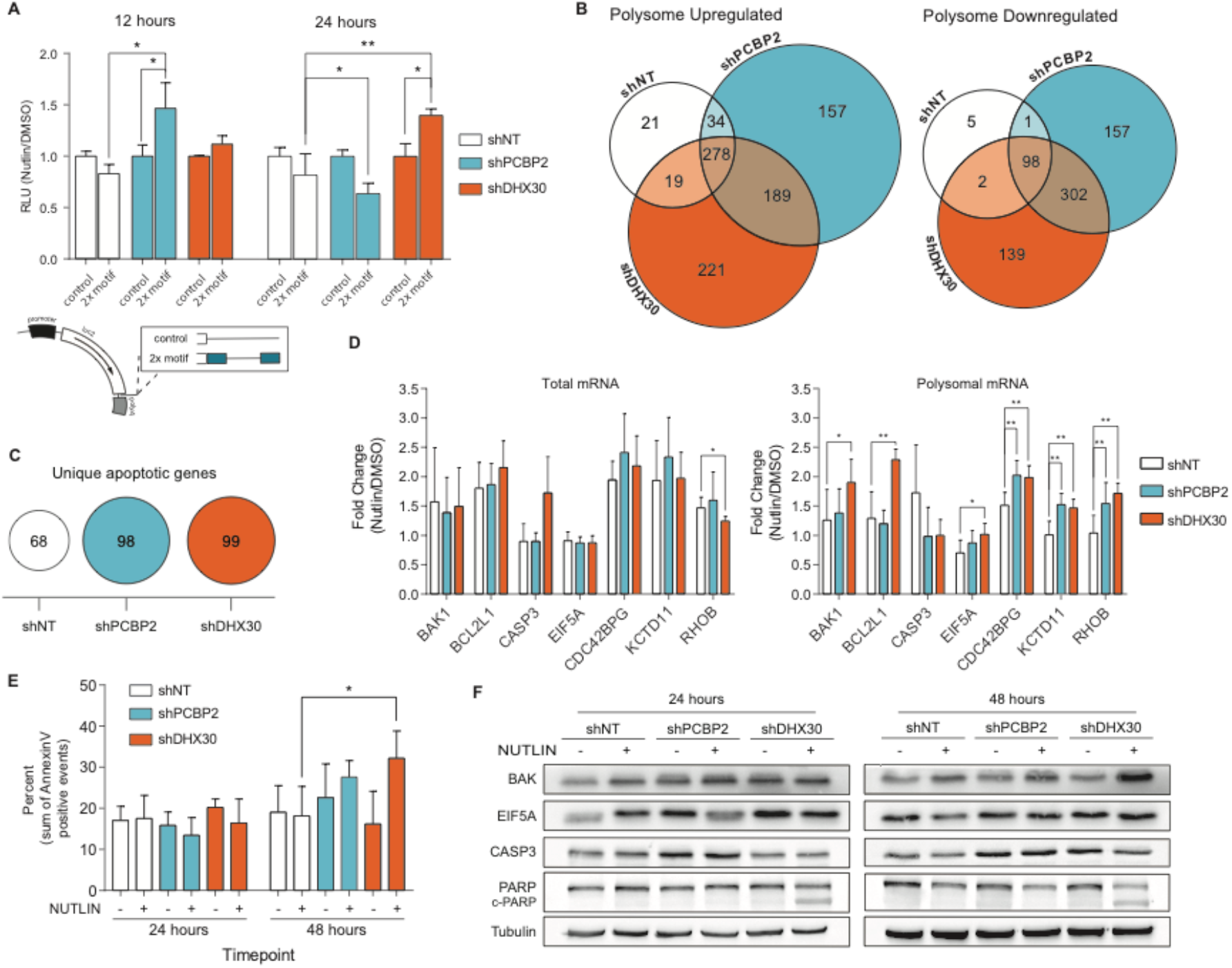
Depletion of PCBP2 and particularly DHX30 in HCT116 cells leads to translational enhancement of motif containing mRNAs and apoptotic cell death in response to Nutlin. **A)** Impact of PCBP2 or DHX30 depletion on the relative translation efficiency of CGPD-motif containing luciferase mRNA. Results were obtained and are presented as in Figure 2D (n=3; * p<0.05, ** p<0.01, Student’s t-test). **B)** Venn diagrams comparing the lists of polysome Upregulated (left) and Downregulated DEGs (right) between HCT116 derivative clones. **C)** Number of unique apoptotic genes among polysomal DEGs (see Tables S6-S8). **D)** Relative RNA expression of selected CGPD-motif genes in total (left panel) and polysomal (right panel) RNA fractions. Presented are the average fold change in HCT116 shNT, shPCBP2 or shDHX30 Nutlin treated cells and the standard deviation of a least four biological replicates. Especially upon DHX30 depletion, BAK1, BCL2L1, RHOB show a tendency to be enriched in the polysomal fraction. (* = p<0.05; ** p<0.01 Student’s t-test) **E)** Results of Annexin-V assay in Nutlin treated HCT116 derivative clones (n=3; * p<0.05, Student’s t-test). **F)** Protein levels for the indicated CGPD-motif genes 24 and 48 hours after Nutlin treatment of HCT116 shNT, shPCBP2 or shDHX30 cells. BAK and EIF5A protein levels are higher in PCBP2 and DHX30 depleted cells. PARP blot is shown as control of apoptotic signaling pathway activation. The cleaved form (indicated as cPARP) is detected upon DHX30 depletion and Nutlin treatment. Relative quantification of the protein levels is presented in Table S15. See also Figure S5.

Next, we compared Nutlin-induced changes in polysomal mRNAs association in HCT116 shNT, shPCBP2, and shDHX30 cells by RNA-seq (**Figure 5B**, **Figure S5B**, **Tables S6**-**S8**). Indeed, depletion of DHX30 or PCBP2 led to a significant increase in the number of DEGs compared to the HCT116 shNT condition (**Figure 5B**, **Figure S5C**). However, we observed a large common core of both upregulated and downregulated DEGs (278 and 98, respectively). Interestingly, a rather large set of genes are in common for shPCBP2 and shDHX30 cells, particularly among repressed DEGs. At the same time, we observed a specific signature for each depleted protein. As expected, upregulated DEGs showed similar enrichment for p53-related ontologies (**Figure S5D**, **Table S9**). Instead, we found a strong enrichment for “DNA repair, replication, S phase, and checkpoint” terms only in shPCBP2 and shDHX30 cells. Using GSEA to better exploit the magnitude of gene expression changes, clear differences among the three HCT116 derivative lines were identified with shDHX30 cell showing stronger enrichment for programmed cell death and p53 downstream pathways (**Figure S5E**, **Tables S10-S12**). Indeed, the number of unique apoptotic genes was about 40% larger in shPCBP2 and shDHX30 cells (**Figure 5C**; **Tables S6-S8**). A *de novo* sequence motif search (Pavesi et al., 2004) was performed on the 5’ and 3’UTRs of polysome associated DEGs. By comparing the newly discovered 3’UTR motifs with the top three CGPD-motifs originally identified in SJSA1, a higher level of correlation was found for the motifs identified in shDHX30 cells with respect to the other models (**Figure S5F**, **Table S13**). Notably, motifs enriched in the 3’UTRs of polysome associated DEGs in shDHX30 cells correspond to a consensus sequence [5’-ATGG(A)CC-3’] that is similar to the central portion of the CGPD-motif identified in SJSA1 translationally enhanced mRNAs. Indeed, high correlation (>0.9, Pearson correlation) was observed between the original CGPD-motif and those enriched in shDHX30 cells. We next examined the relative changes in polysome association of the 193 CGPD-motif-containing mRNAs previously identified as translationally upregulated in SJSA1 cells (**Figure S2**). An enrichment in Nutlin-induced polysome association was apparent particularly for shDHX30 cells (**Figure S5G**). Eventually, we validated seven genes derived from both the original CGPD-motif list and the newly identified, correlated motif by RT-qPCR and confirmed a significant increase only in the polysomal fractions, particularly in HCT116 shDHX30 cells (**Figure 5D**).

These results suggest that during Nutlin treatment, especially DHX30 silencing may reshape the HCT116 translatome towards the expression of CGPD-motif genes.

### DHX30 depletion enhances Nutlin-dependent apoptosis in HCT116 cells

We then asked whether DHX30 depletion could affect the response of HCT116 cells to Nutlin. HCT116 shDHX30 cells exhibited a higher proportion of cell death based on annexin-V and propidium iodide staining, at 48 hours after Nutlin treatment (**Figure 5E**). Indeed, while shNT HCT116 cells have an apoptotic level that does not exceed 20% as previously reported (Tovar et al., 2006), in shDHX30 this level reaches 30%.

Consistently, the cleaved form of the PARP protein showed a significant increase in Nutlin-treated shDHX30 cells as revealed by western blot (**Figure 5F**). BAK, EIF5A, and CASP3 - examples of CGPD-motif containing genes- were also examined at the protein level (**Figure 5F**- **Table S15**). Consistent with the results in SJSA1 cells, western blot analyses showed relative increases in BAK and EIF5A protein levels more evident 48 hours after Nutlin treatment, particularly in shDHX30 cells. Also consistent with results in SJSA1 (Figure 2C), EIF5A protein showed an increase in shNT cells 24 hours after treatment, but a reduction at the 48 hour time point that is attenuated in shPCBP2 (**Figure 5F).**

Overall, we show that, by modulating DHX30 expression, we can shift the outcome towards apoptosis in HCT116 cells.

### DHX30 depletion increases the sensitivity to Nutlin of U2OS cells

To begin exploring the general role of DHX30 in translation control and modulation of cell death downstream of p53 activation, we employed U2OS cells. Similarly to SJSA1 these cells are p53 wild-type and derived from an osteosarcoma, but in contrast to SJSA1, they undergo cell cycle arrest in response to Nutlin treatment (Tovar et al., 2006). Interestingly, U2OS cells express higher levels of DHX30 compared to SJSA1 (**Figure 6A**). We then tested whether U2OS cells behave similarly to HCT116 cells upon DHX30 depletion. As observed for shDHX30 HCT116 cells, U2OS are more sensitive to Nutlin upon DHX30 depletion, particularly at 72-hours, based on the survival rate and on the proportion of PI-positive cells (**Figure 6B**). However, we did not observe an appreciable increase in the fraction of annexin-V positive cells (data not shown). Next, by employing the luciferase reporter assay based on the cloned CGPD-consensus motif we observed that the impact of the motif already apparent in shNT cells is slightly increased in shDHX30 cells (**Figure 6C**). Total and polysomal RNAs were extracted from U2OS shNT and and shDHX30 treated with Nutlin for 24 hours. DHX30 depletion led to higher expression as well as higher polysome association of three CGPD-motif mRNAs tested by qPCR (**Figure 6D**). Contrary to what we observed in HCT116 cells, DHX30 depletion *per se* resulted in higher expression of EIF5A and CASP3. The depletion of DHX30 resulted in higher constitutive protein levels for CASP3, BAK, and EIF5A (**Figure 6E**). The induction of BAK expression in response to Nutlin was consistent among qPCR and western blot experiments, as highlighted in **Table S15**. These data suggest that, also in U2OS, DHX30 contributes to define the cell outcome possibly modulating the expression of CGPD-motif genes.

**Figure 6.**
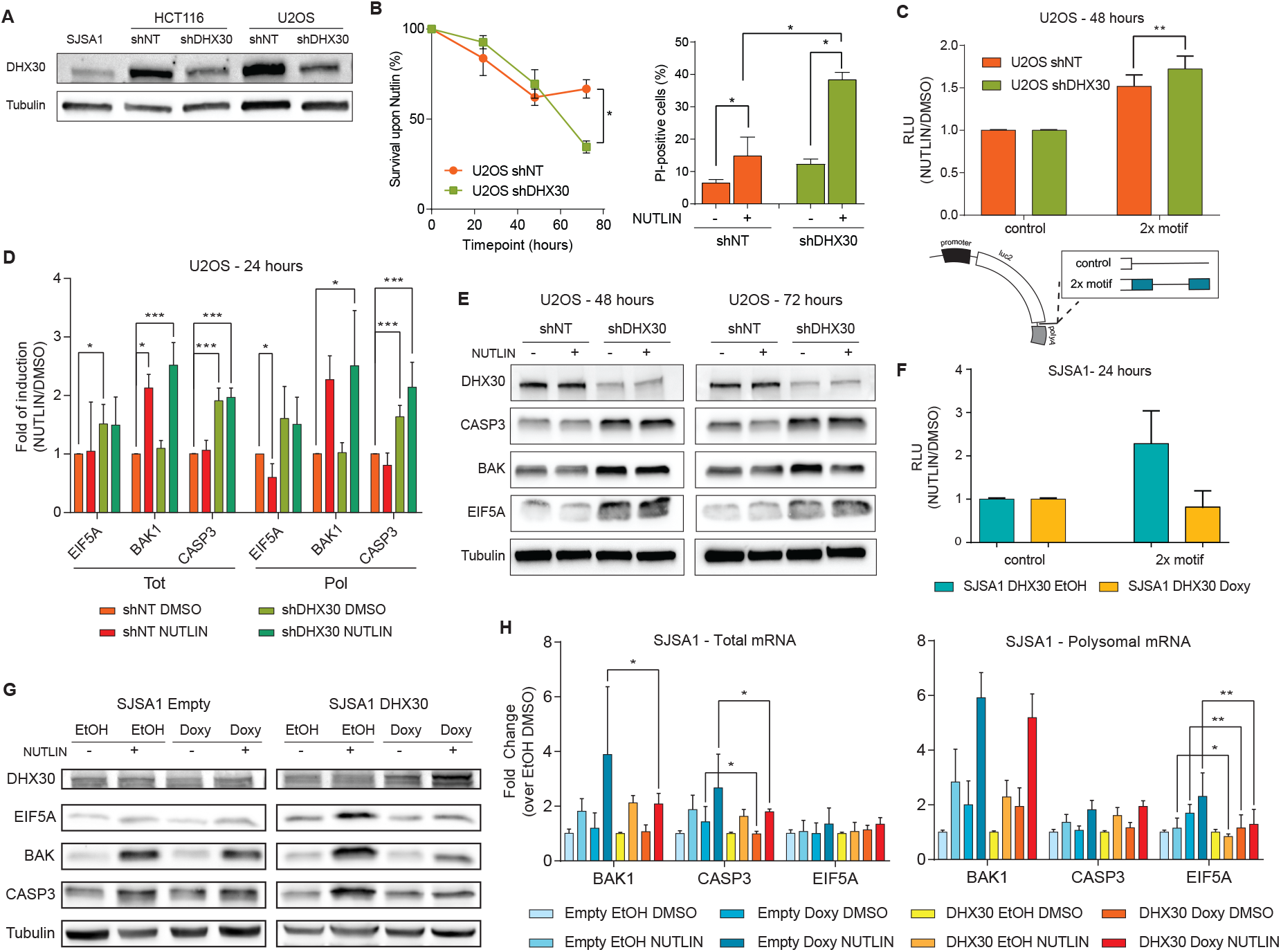
Depletion of DHX30 in U2OS results in higher EIF5A, BAK, CASP3 expression and enhanced sensitivity to Nutlin. **A)** Relative DHX30 protein expression levels in SJSA1, HCT116 shNT or shDHX30, and U2OS shNT or shDHX30. U2OS and SJSA1 are both osteosarcoma cell lines with a different cell fate upon Nutlin treatment, as previously reported. **B)** Left: survival curves in response to Nutlin treatment were obtained by high-content imaging. Right: percentage of cells permeable to Propidium Iodide, used as a marker of cell death, were also measured using high-content imaging. Presented are the average and standard deviations of three replicates (*, p<0.05, Student’s t-test). **C)** Reporter assay with the construct containing two copies of the consensus CGPD-motif inserted in the β-globin 3’UTR placed downstream of the luciferase gene. U2OS cells depleted for DHX30 or control were transfected with the plasmids and treated with Nutlin 24 hour after transfection. The average ratios of luciferase activity normalized on the renilla activity and on the luciferase RNA values are presented along with the standard deviation of three biological replicates (**. p<0.01, Student’s t-test). **D)** Impact of DHX30 depletion or/and Nutlin treatment on the relative expression of the CGPD-motif mRNAs EIF5A, BAK1, and CASP3 in total or polysomal RNA extracts. Bars plot the average fold change relative to the DMSO control for U2OS shNT cells. The standard deviation of four biological replicates are presented. (*, p<0.05; **, p<0.01; *** p<0.001; Student’s t-test). **E)** Western blot of selected CGPD-motif containing targets (EIF5A, BAK, CASP3) at two time points (48 and 72 hours) after Nutlin treatment. The 72 hour time point was chosen based on the results from the survival curves. **F)** The luciferase assay with the constructs containing two copies of the consensus CGPD-motif or its control was performed in SJSA1-DHX30 cells, exposed to 2.5μg/ml of doxycycline (or EtOH) for 24 hours prior to the treatment with Nutlin for 24 hours. As done for panel C, the normalized average ratios and standard deviations of three biological replicates are presented. **G)** Representative example of a western blot for DHX30, EIF5A, BAK and CASP3 in SJSA1-Empty or SJSA1-DHX30 cells exposed to doxycycline for 24 hours and then to Nutlin for 48 hours. Quantification of immunoblots at different time points are presented in Figure S6 and Table S15. **H)** Impact of DHX30 overexpression on the relative levels of BAK1, CASP3, and EIF5A, in total (left panel) or polysomal (right panel) mRNAs. Bars plot the average fold change relative to EtOH-treated DMSO controls for SJSA1-Empty or SJSA1-DHX30. Cells were exposed to 2.5μg/ml of doxycycline (or EtOH) for 24 hours prior to the treatment with Nutlin (or DMSO) for an additional 12 hours. The standard deviations of at least two biological replicates are presented. (*, p<0.05; **, p<0.01; *** p<0.001; Student’s t-test). Results for BCL2L1, p21, and DHX30 are presented in Figure S6.

### DHX30 overexpression in SJSA1 cells reduces the translation of CGPD-motif mRNAs

Finally, we established an inducible overexpression system for DHX30 in SJSA1 cells as a proof-of-principle of the effect of this protein on the expression of CGPD-motif mRNAs. Exposure to doxycycline led to modest upregulation of DHX30 (**Figure 6H**, **Figure S6**). As expected DHX30 overexpression was associated with a reduction in the activity of the CGPD luciferase reporter (**Figure 6F**). Comparing SJSA1-Empty with SJSA1-DHX30 cells we observed a reduction of EIF5A, BAK, and CASP3 protein levels (**Figure 6G**), that was associated with lower amounts of BAK1 and CASP3 total mRNAs and reduced polysomal association of EIF5A mRNA as a consequence of DHX30 overexpression (**Figure 6H**). We have thus described here a novel post-transcriptional regulatory mechanism that reprograms the translatome to affect the apoptotic choice and thus the phenotypic outcome of Nutlin treatment.

## DISCUSSION

To analyze gene expression changes associated with the commitment to p53-dependent apoptosis, we deliberately chose to exploit the expected difference among cell line models derived from different tissue types. To obtain robust and specific activation of p53 without the overt engagement of DNA damage responses, we challenged them with the MDM2 inhibitor Nutlin-3a. We reasoned that the heterogeneity of these cell lines would enable us both to characterize if a common gene expression program is consistently activated by p53 and to identify if changes in translational efficiency can better explain distinct phenotypic commitments. Our data demonstrate that p53 activates a conserved, multifunctional transcriptional program in HCT116 and SJSA1 cells, regardless of the response to Nutlin (**Figure 1**) (Andrysik et al., 2017). However, cellular responses progressively diverge from p53 promoter targeting to protein synthesis and are the most distinct at the translational level, while still being coherent with the cell fate choices. Indeed, translationally-enhanced mRNAs captured from SJSA1 cells 12 hours after Nutlin treatment - a time point that precedes their apoptosis engagement - are enriched for modulators of programmed cell death (**Figure 1**, **Figure S1**). In a non-mutually exclusive view, these results may suggest that the initial p53-dependent response to Nutlin is, in large part, dependent on the transcriptional core program. This transcriptional response then builds-up a specific post-transcriptional program, which regulates translational efficiency through the functional interaction of mRNA motifs and trans-factors.

Indeed, by scanning the UTRs of SJSA1 translated mRNAs, compared to all other lists of differentially expressed genes from the two cell lines, we discovered one specific 3’-UTR cis-element. We showed this CGPD-motif to be sufficient to confer cell-line-, treatment- and p53-dependent stimulation of translation. This *cis*-element led us to identify RNA binding proteins that are either common or specific binders from HCT116 and SJSA1 cell extracts. PCBP2 was identified as interactor in both cell lines, while DHX30 was specific for HCT116 cells.

Poly(C)-binding proteins (PCBPs) are a family of five RBPs characterized by high-affinity, sequence-specific interaction with poly(C). The most widely studied PCBPs are hnRNP K, αCP-1, and αCP-2, the latter also known as PCBP1 and PCBP2 or hnRNP E1 and hnRNP E2, respectively (Kiledjian et al., 1995). Recently, two other members of the PCBP family were discovered and named PCBP3 and PCBP4 (Makeyev and Liebhaber, 2000). All members of this family are evolutionarily related, and the common feature of all PCBPs is the presence of three hnRNP K homology domains (Makeyev and Liebhaber, 2000). Members of the PCBP family act as mRNA stabilization factors through interactions with the 5′-untranslated regions (Holcik and Liebhaber, 1997) or as translational silencers by binding to 3′-untranslated regions (Collier et al., 1998; Ostareck et al., 1997). PCBP2 was recently linked to the regulation of p73 expression, antioxidant response, neural apoptosis but also glioma cell growth, apoptosis, and migration (Lin et al., 2016; Mao et al., 2016; Ren et al., 2016).

DHX30 is a poorly characterized, evolutionarily conserved protein containing a DExH helicase motif, whose ortholog in mice was shown to be essential for embryo development (Zheng et al., 2015). In human cells, DHX30 appears to play a role in antiviral responses (Zhou et al., 2008). The protein can also localize to the mitochondria where it may play a role in mitoribosome biogenesis (Antonicka and Shoubridge, 2015). Notably, DHX30 has been recently associated with translation regulation of human mRNAs; it has indeed been identified as a component of the mammalian ribo-interactome (Simsek et al., 2017). Moreover, specific mutations were shown to impair its RNA recognition or ATPase activity and, when overexpressed, led to stress granule formation and inhibition of global translation (Lessel et al., 2017).

Considering amino acid sequence and functional motifs, DHX30 has four main paralogs in human, with protein identity ranging from nearly 30% to 20%. The closest one is DHX36, which has been involved in the control of mRNA stability (Bourgeois et al., 2016). The most studied paralog is DHX9, a protein that is primarily localized in the nucleus, where it impacts various processes including transcription, splicing, and RNA export (Bourgeois et al., 2016). Recently, the dependency of cancer cells on higher levels of DHX9 has been proposed and in part associated to p53 wild-type activity (Lee et al., 2016). Interestingly, DHX9 silencing in U2OS cells led to cell cycle arrest.

Related to the mRNA fate of p53 target genes, PCBP2, PCBP1, and PCBP4 have been reported as inhibitors of p21 mRNA translation (Scoumanne et al., 2010). Using multiple approaches, we confirmed the interaction between PCBP2 and p21 mRNA in HCT116 cells. We discovered that DHX30 can also bind the p21 mRNA in HCT116 cells (**Figure 4D**). Overall, p21, EIF5A and BAK1 are just examples among several PCBP2 and DHX30 shared targets, as confirmed by our reanalysis of publicly available eCLIP data (Van Nostrand et al., 2016). Even if we cannot completely rule out that other PCBP-DHX family proteins also participate in the translational modulation of mRNAs through the CGPD-motif, our data strongly support the view that PCBP2 and DHX30 bind the CGPD-motif, and that their cell-type specific interaction, possibly resulting on the differences in relative expression (**Figure S3C**), may contribute to the observed outcome.

Here, we focused on elucidating the role of DHX30 and PCBP2 in shaping the translatome, and thus, the observed phenotype, upon p53 activation.

Polysome profiling of HCT116 depleted for PCBP2 or DHX30 revealed that the number of mRNAs changing their polysomal association in response to the treatment with Nutlin is much larger compared to control cells (**Figure 5B**). Gene Ontology and Gene Set Enrichment Analysis revealed significantly enriched terms consistent with p53 activation with a higher number of apoptotic targets being modulated in the depleted PCBP2 and DHX30 depleted cells consistent with activation of apoptotic responses, particularly in HCT116 shDHX30 cells. Notably, the 3’UTRs of polysome upregulated mRNAs were enriched for a motif that correlates with the CGPD-motif originally found in the SJSA1 translatome (**Figure S5F**). Importantly, the expression levels of the CGPD-motif genes increases upon DHX30 depletion and Nutlin treatment (**Figure 5D**, **Figure S5G**). Our data strongly show that DHX30 depletion can recapitulate features of the SJSA1 translatome. Consistently, starting from U2OS cells, an osteosarcoma cell line that expresses wild-type p53 but that undergoes Nutlin-dependent cell cycle arrest, we observed that DHX30 depletion led to increased Nutlin sensitivity, associated with enhanced protein expression of CGPD-motif targets (**Figure 6**). In addition, moderate DHX30 overexpression in SJSA1 cells led to reduced translation of CGPD-motif mRNAs. Unexpectedly, we noticed a synergistic interaction between doxycycline and Nutlin in the induction of DHX30 mRNA and protein (**Figure S6**).

Depletion of PCBP2 in HCT116 cells resulted in less pronounced changes in mRNA translation, a weaker correlation between an over-represented 3’UTR motif of translated DEGs and the CGPD-motif, but similar proportions of predicted PCBP2/DHX30 targets. Along with the results from motif pull-down and RIP experiments in depleted cells as well as with co-immunoprecipitation studies, our data indicate a functional cooperation between PCBP2 and DHX30, whereby abolishing PCBP2 binding strongly reduces DHX30 interaction to the CGPD-motif as well (**Figure 4**). Our data do not demonstrate if there is a direct interaction between the two proteins. Nevertheless, we can suggest that their interaction is RNA-dependent. We thus propose that both DHX30 and PCBP2 can bind RNA and shared RNA substrates would tether them together. Hence, depletion of one protein could impact on RNA binding of the other also through an indirect effect on the target RNA structure. It is likely that other proteins are involved along with PCBP2 and DHX30 in the interaction with target RNAs at CGPD-motif sites, as suggested also by the mass-spectrometry data. As observed for other families of RBPs, the eventual mRNA fate can depend on the results of the interplay between different RBPs or RBP complexes targeting the same specific cis-element (Dassi, 2017; Lal et al., 2004). This hypothesis is supported by the fact that DHX30 depletion led to a significant increase in Nutlin-dependent apoptosis 48 hours after treatment, while PCBP2 silenced cells did not exhibit more Nutlin-dependent apoptosis (**Figure 5E, 5F**). Hence, it appears that DHX30 could act as a limiting factor for the apoptotic commitment of cells downstream of p53 activation by Nutlin. From a broader perspective, these results envision the potential clinical benefit of targeting DHX30 in cell-cycle arrested cells. The relative expression levels of DHX30 could also be instrumental in understanding which cancer cells to target. Even though PCBP2 and DHX30 are not direct p53 targets according to ChIP-seq data of Nutlin treated cells (Allen et al., 2014; Nguyen et al., 2018), future studies may reveal the possibility that p53 indirectly controls PCBP2 and DHX30 interaction with each other and with their shared target RNA motif we have discovered.

Our work highlights the relevance of translational specificity and efficiency as a largely unexplored regulatory layer in the commitment to programmed cell death downstream of p53 activation. We described in detail one of the many possible post-transcriptional regulatory mechanisms implicated in the apoptotic outcome, which could potentially lead to new avenues for predicting treatment outcome and devising more effective intervention strategies.

## Supporting information

Supplemental Figures and List of Supplemental Tables

## ACKNOWLEDGMENTS

We wish to thank Veronica De Sanctis and Roberto Bertorelli -CIBIO NGS CIBIO core facility- and Isabella Pesce -CIBIO Cell Analysis and Separation core facility- for technical support in RNA-sequencing and apoptosis assays. We wish to thank Laura Pezzè, Bartolomeo Bosco and Natthakan Thongon for technical assistance.

## FUNDING

This work was supported by the Italian Association for Cancer Research, AIRC (grants IG# 12869 and #18985 to A.I.) and by NIH grant 2R01CA117907. Sara Zaccara was supported by an AIRC Reintegration Fellowship.

## AUTHORS’ CONTRIBUTIONS

All Authors have read and agreed on this submission and declare no competing financial interests. S.Z., D.R., A.R. J.M.E, and A.I. conceived experiments; S.Z. E.D. M.D.G., D.R., A.R., Z.A., K.D.S. performed experiments; E.D., S.Z., M.D.G., D.R. A.R. analyzed data; S.Z. and A.I. wrote the first draft; S.Z., M.D.G., E.D., D.R., A.R., J.M.E., A.Q, and A.I. revised the manuscript; J.M.E. and A.I. secured funding; J.M.E. and A.I. supervised the study.

## DECLARATION OF INTERESTS

“The authors declare no competing interests.

## ACCESSION NUMBERS

RNA-sequencing data are deposited in GEO: for HCT116 shNT, shPCBP2 and shDHX30 cells (GSE95024); for parental HCT116 and SJSA1 cells (GSE86222).

## MATERIALS AND METHODS

### Cell lines and culture condition

HCT116, SJSA1 were maintained in RPMI (Corning); U2OS cells were maintained in DMEM (Corning). Media were supplemented with 10% FBS, antibiotics (100 units/mL penicillin plus 100 mg/mL streptomycin) and 2mM L-glutamine. SJSA1 p53KO cells were provided by the Espinosa lab. All cell lines were tested for mycoplasma contamination and authenticated by STR profiling (BMR Genomics). When needed cells at 70-80% of confluence were treated with 10μM Nutlin (Sigma-Aldrich) dissolved in DMSO.

### Polysome Profiling and RNA extraction

Cells were allowed to reach 60-70% confluence before treatment with 10μM Nutlin. After 12 hours, polysomal separation was performed as previously described (Dassi et al., 2015; Provenzani et al., 2006; Zaccara et al., 2014). Briefly, samples were loaded in 15-50% linear sucrose gradients, ultra-centrifuged and fractionated with an automated fraction collector. All the fractions containing subpolysomal (fractions up to the 80S density) or polysomal RNA (fractions containing two or more polysomes) were identified and pooled in two separate tubes. RNA was purified by extraction with 1 volume of phenol-chloroform and, after isopropanol precipitation, a washing step in 70% v/v ethanol was performed in order to remove phenol contaminations. Two biological samples were analyzed for SJSA1 and HCT116 parental cells. Four biological replicates were analyzed for HCT116 derivative clones that were depleted for PCBP2 or DHX30 and for U2OS shNT control and U2OS shDHX30. RNAs were extracted using acidic Phenol-Chloroform, as previously described (Zaccara et al., 2014). Polysome profiling was also performed with U2OS with the same protocol but after a 24-hour Nutlin treatment.

### DHX30 and PCBP2 silencing and CRISPR/Cas9 based knock-out

HCT116 cells were transduced with lentiviral vectors containing a pLK0.1 plasmid expressing shRNA sequences against DHX30 or PCBP2. Plasmids were obtained by the Functional Genomics Facility (http://functionalgenomicsfacility.org/shRNA). Sequences for DHX30 are: TRCN0000052028 (GCACACAAATGGACCGAAGAA) TRCN0000052031 (CCGATG GCTGACGTATTTCAT), TRCN0000052032 (GAGTTGTTTGACGCAGCCAAA), Sequences for PCBP2 are: TRCN0000074685 (GCCATCACTATTGCTGGCATT) and TRCN0000074687 (CCTGGCTCAATATCTAATCAA). A scramble sequence SHC206 was also used. In order to obtain a reproducible and efficient transduction, 1) cells were transduced with the same amount of lentiviral particles previously quantified for their RT units (Barczak et al., 2015), 2) cells were spin-inoculated for 2 hours at 1600 RCF. The day after, lentivirus containing media was substituted with fresh media. After 48 hours, cells were collected for a preliminary screening and selection of transduced cells was performed by 0.5 μg/mL puromycin. Single cell clone selection was then performed. At least one clone for each shRNA was obtained (data not shown). To facilitate our experiments, the clone showing the best silencing effect was chosen for further analysis including polysomal profiling and RNAseq analysis. Puromycin (Sigma-Aldrich) was used to maintain the selection, at 0.1 μg/mL as final concentration.

As alternative strategy, we attempted to obtain complete DHX30 knock-out clones using CRISPR/Cas9. Two different guides (G1: CTAGTCTACGTGCACACAAATGG; targeting exon 6 of ENST00000348968.8 or ENST00000457607.1 or G2: CATAAAATGGCCCAAGAGCGTGG targeting exon 7) were cloned in a modified pX330 vector that also contains puromycin selection marker. Plasmids were transfected in HCT116 and transformants were selected by puromycin. The efficiency of generating indel at the expected Sp-Cas9 cut site was monitored by PCR amplification with specific primers (Table S14), Sanger sequencing, and TIDE (Brinkman et al., 2014). Single clone isolates were then obtained by limiting dilution. Despite high efficiency of indel generation, no complete knock-out clones were obtained (Figure S4A), suggesting that DHX30 is essential in HCT116. Anecdotally, DHX30 is considered essential in 80/625 cell lines based on CRISPR screening in the DepMap portal of the Achilles project (Ghandi et al., 2019), including cancer cell lines of intestinal origin. However, we did not find data on the essential nature of the DHX30 gene in HCT116 cells. PCBP2 is instead classified as a common essential gene in DepMap and we did not pursue CRISPR knock-out.

### DHX30 inducible overexpression in SJSA1 cells

Full length wild type human DHX30 cDNA corresponding to transcript (ENST00000348968.8) was cloned into the TetON lentiviral vector pCW57.1 (Addgene), exploiting the Nhe I and Age I restriction endonucleases. Stable SJSA1-Empty (transduced with the empty pCW57.1 vector) and SJSA1-DHX30 transformant cells were selected exploiting the puromycin marker. Conditions for inducible overexpression were established using different concentrations of doxycycline, choosing 2.5μg/ml as the optimal dose.

### Library preparation and data analysis for HCT116 and SJSA1 datasets

RNA concentration, purity and integrity were measured by the Agilent 2100 Bioanalyzer (Agilent Technologies) discarding RNA preparations with RIN (RNA integrity number) value <8. PolyA+ mRNA isolation was performed using the Dynabeads mRNA DIRECT Micro kit (Life Technologies) following the “mRNA isolation from purified total RNA” protocol and using 1.5 μg of RNA as input. Then, we proceeded to the library preparation according to the Ion Total RNA-Seq Kit instructions. As indicated, we assessed the quality and efficiency of each preparation step using the Agilent RNA 6000 Pico kit with Agilent 2100 Bioanalyzer instrument. Subsequently, each library template was clonally amplified on Ion Sphere Particles for sequencing on the Ion Proton System, producing ≈60-80 M raw reads per sample. After quality filtering and trimming (minimum read length: 30nt; maximum read length: 150nt) by the FASTX-Toolkit (http://hannonlab.cshl.edu/fastx_toolkit), mapping to the hg19 build of the human genome (February 2009 GRCh37, NCBI Build 37.1) was performed using GSNAP (Wu and Nacu, 2010). Mapping parameters allowed for up to 3% mismatches per read to avoid bias against alignment of longer reads. To compute the per-gene read counts, we used HTSeq (Anders et al., 2015). We chose the intersection non-empty mode for reads overlapping more than one gene/exon feature. The DESeq R package (Anders and Huber, 2010; Love et al., 2014) was used to call Differentially Expressed Genes (DEGs) starting from two replicates for each condition and comparing each fraction with itself across conditions (*e.g.* Nutlin total versus DMSO total; Nutlin polysomal vs DMSO polysomal). For all analyses on DEGs, two thresholds were set: (1) log_2_(fold-change) >1 and <1 for up-regulated and down-regulated genes, respectively; (2) FDR-corrected p-value <0.1. We defined coupled DEGs as genes which met the indicated thresholds at the total and polysomal RNA level. Translationally enhanced or reduced genes are genes that met the indicated thresholds in the polysomal fraction, but are not significant at the total RNA level. Un-changed in translation DEGs are genes that follow the reported thresholds in the sub-polysomal fraction, but not in the polysomal one independently from their expression change at the total RNA level.

### Library preparation and data analysis for shDHX30, shPCBP2 and scramble datasets

Sequencing libraries were constructed following the TruSeq RNA Library preparation kit v2 manufacturer instruction and using 1.5 μg of RNA as input. We assessed the quality of our input RNA using the Agilent RNA 6000 Nano kit with Agilent 2100 Bioanalyzer instrument. Each condition was analyzed using four replicates. A total of 24 samples were sequenced using HiSeq 2500, producing ≈25-28 M raw reads per sample. After quality filtering and trimming with trimmomatic (minimum quality 30, minimum length 36nt) (Bolger et al., 2014) we quantified each Gencode v27 (http://www.gencodegenes.org/releases/) transcript by means of Salmon (Patro et al., 2016). edgeR (Robinson et al., 2010) was used to call Differentially Expressed Genes (DEGs), considering a 0.05 significance threshold on the adjusted p-value.

### Enrichment, Gene Ontology and Pathway analysis

Pathways and gene ontology enrichment analysis for all our selected categories of coupled or uncoupled DEGs were performed with Metascape (http://metascape.org) (Tripathi et al., 2015). We used the multiple lists input format and a custom analysis selecting Reactome Gene Sets, Canonical Pathways, GO Biological Processes, and Hallmark Gene Sets. Enrichment analyses were performed by means of the Fisher test, with a significance threshold of 0.05 on the enrichment p-value, and a background gene set consisting of all Gencode v25 genes (http://www.gencodegenes.org/releases/). GSEA was performed with the fgsea R package (Sergushichev, 2016), using the hallmark, canonical pathways, and GO gene sets, a significance p-value threshold of 0.05 and 10000 permutations to compute the p-value. Apoptotic genes were defined as those annotated with the apoptotic process GO term (GO:0006915).

### Motif discovery

To search for common sequence motifs in each category of coupled or uncoupled DEGs, we used Weeder (Pavesi et al., 2004), setting the following parameters: (1) longest 5’ or 3’UTR of each gene as input, (2) motifs had to be found in at least 25% of all input sequences, (3) motif length ranging from 6 to 12nt, (4) motif search performed on the sense strand only. Background for Weeder consisted in the frequency of all possible n-mers (with n=6,8,10,12) in the human genome, as provided by the Weeder package. Best motifs were selected by the “adviser” program of the Weeder suite, especially according to their redundancy. Resulting motifs were compared by computing the Pearson correlation of their positional weight matrices by means of the TFBStools R package (Tan and Lenhard, 2016). Genome-wide occurrences of interesting motifs were searched with FIMO (Grant et al., 2011), using the motif matrix derived by Weeder and a threshold of 1e-04 on the match p-value.

### Cloning strategy and luciferase assay

Full-length β-globin 3’-UTR, β-globin 3’-UTR with the addition of a C-rich motif at the 3’UTR beginning or at the 3’UTR end or both, were cloned after the luciferase stop codon in the pGL4.13 vector exploiting the XbaI restriction site. Primers used for the cloning are reported in Table S14. Dual-luciferase reporter assay was done according to the manufacturer’s instructions (Promega). Briefly, cells were plated on a 24-well plate, and then transfected with control pGL4.13 reporter vector or the same vector containing the different 3’UTRs, and a Renilla luciferase vector. After 24 hours, cells were treated with DMSO or Nutlin. Luciferase activity was measured in triplicate using Infinite 200 PRO microplate reader (TECAN) 12, 24 or 48 hours post-treatment. Firefly values were normalized on the Renilla (RLU). To highlight changes upon treatment, data were then normalized on the DMSO condition. From the same samples, RNA was also extracted by TRIzol reagent (Thermo Fisher scientific) in order to quantify the luciferase mRNA levels. The mRNA levels were then used to normalize the luciferase values.

### Western Blotting

Antibodies used for Western Blot analysis were p53 (DO-1), β-Tubulin (3F3-G2), hnRNP-E2 (23G), EIF5A (H8), Bak (G-23) from Santa Cruz Biotechnology; DHX30 (ab85687), PCBP1 (ab154252), MCG10 (ab59534), p21 (ab1676), PABP (ab21060) from Abcam;, MYH9 (A304-490a) from Bethyl, CASP3 (gtx110543) from GeneTex. Cells were seeded and allowed to reach 70-80% of confluence before Nutlin treatment for 24, 48 or 72 hours. Proteins were extracted using RIPA buffer, supplemented with protease inhibitors (Sigma-Aldrich) as previously described (Zaccara et al., 2014) and quantified using the BCA assay (Thermo Fisher Scientific). The relative molecular mass of the immunoreactive bands was determined using PageRuler Plus Prestained Protein Ladder (Thermo Fisher Scientific). The semi-quantitative analysis was performed using GAPDH or Tubulin as reference proteins for loading control.

### Reverse transcription and qPCR

cDNA was generated from 0.5-2 μg of RNA using the RevertAid First Strand cDNA Synthesis Kit (Thermo Fisher Scientific) in 20 μL final volume following manufacturer’s instructions. All qPCR assays were performed on a CFX Touch Real-Time PCR Detection System (Bio-rad,). We validated targets using the 2X KAPA SYBR® FAST qPCR Kit (Kapa Biosystems, Resnova) or qPCR Biosystem SYGREEN separate ROX mastermix (Resnova), 0.2 μM of Forward and 0.2 μM of Reverse primers purchased from Eurofins Genomics (Eurofins Genomics, Germany GmbH) and 20-25ng of cDNA. The list of primers is presented in Table S14. We present the mRNA quantification relative to the DMSO condition for each fraction (tot, pol) in order to highlight changes upon treatment. The relative quantification was obtained using the comparative Cq method (ΔΔCq), where GAPDH or YWHAZ served as reference genes. The relative folds of change were analyzed using t-test considering at least two independents biological replicates, as indicated in the various figures.

### RNA pull-down

Two in vitro synthesized RNA probes with a 5’-UTR biotinylation were purchased from IDT (Integrated DNA Technologies). Their sequence match the wild-type motif sequence (wt: 5’-GAAGGGCCCU CCCCAUGGCCCU GGAGAGUGGG-3’), or the mutant one (mut: 5’-GAAGGGCCCU CCCCAUGGAGAU GGAGAGUGGG-3’). These probes were used to perform pull-down experiment as previously reported (Dassi et al., 2013). Briefly, after washing and immobilization of 0.50mg/reaction of streptavidin-conjugated beads (M-280 – Thermo Fisher Scientific) according to the manufacturer’s instruction, the binding of 80 pmol RNA probes to beads was obtained by incubation for 20 minutes at room temperature using gentle rotation. After washing, 500 μg of protein extract was added and incubated for 1h at 4°C to the streptavidin beads now binding the biotinylated RNA probe. After washings, the bound complex was detached from the beads heating at 95°C for 7 minutes and then load on a 8% polyacrylamide gel. In case of mass-spectrometry analysis the gel was stained by Coomassie. The visual comparison of bands from the SJSA1 and HCT116 cell pull-down upon DMSO or Nutlin treatment was used to select a range of protein size for quantitative mass spectrometry that showed the most evident difference between the two cell lines. Label-free quantification was performed at the IFOM Proteomics facility (Milan) (Table S5). We filtered the list of obtained proteins according to two parameters: i) number of matching peptides greater than 2; ii) sequence coverage greater than 10%. In case of DHX30 and PCBP2 detection, classical protein transfer and antibodies incubation procedures were applied. Input samples (30 μg) were loaded as control.

### Co-Immunoprecipitation

Magnetic Protein G or A beads were washed twice with 200ul of CHAPS buffer. Then, beads were resuspended in 200μl of CHAPS buffer in the presence protease inhibitors RNase inhibitors- and 3μg of PCBP2 antibody. Incubation was performed at 4°C for 1h to let the antibody attach to the beads. beads-conjugated antibodies were washed with 1 mL CHAPS buffer to discard excess of unbound antibody. 1 mg of cell lysate previously obtained using CHAPS buffer -in the presence of protease, and RNase inhibitors - and gently rotating at 4°C for 30 minutes was added to the beads-conjugated antibodies. We performed overnight incubation at 4°C favoring gentle rotation of the samples. To test whether the binding of the antibody was RNA-dependent, before the incubation, one of the sample lysates was also pre-treated with 10 μg/mL RNase A for 10 minutes at 37 °C. Three washes with CHAPS buffer rotating the samples for 5 minutes at 4°C were performed. The complex was then resuspended in 30μl of loading buffer with 2% Urea and 20% DTT. After denaturation at 95°C for 8 minutes, the eluate was loaded on the 8% acrylamide gel. As comparison, 30μg of input cell lysate were also loaded using the same buffer conditions. DHX30 and PCBP2 were detected using classical protein transfer and antibodies incubation procedures. PABPC1 was used as positive control of PCBP2 immunoprecipitation (Wan et al., 2015).

### RIP assays

∼10^7^ cells were lysed in 1mL of Lysis buffer (100mM KCl, 5mM MgCl2, 10mM HEPES pH7, 0.5% NP-40, 1mM DTT, 1U/ul RNase Inhibitors, 1X Protease Inhibitor Cocktail) using a scraper. Lysates were transferred in a falcon tube, left at least 2 hours at −80°C, centrifuged at 10000 rpm for 30 minutes and supernatants were collected in a new tube. Dynabeads ProteinA or ProteinG (Thermo Fisher scientific, depending on the antibody species) were prepared by washing them twice with NT2 Buffer (50mM Tris-HCl pH7.4, 150mM NaCl, 1mM MgCl_2_; 0.05% NP40) and resuspended in NT2 buffer. Beads were distributed in different tubes, supplemented with twice their initial volume of NT2 Buffer. The specific antibody (3 μg for PCBP2 or 5 μg for DHX30) or IgGs were added to the beads and incubated for 2 hours on a wheel at 4°C. Lysates were pre-cleared by adding a mix containing ProteinA and ProteinG Dynabeads in equal amount and left for 1 hour at 4°C on a wheel. After placing the tubes on a magnet, supernatants were collected and 1% of their volume was used as input to be directly extracted with TRIzol. The remaining supernatant was added to the antibody-coated beads and incubated overnight on a wheel at 4°C. Beads were resuspended in 1ml NT2 buffer, transferred in a new tube and washed with 1ml NT2 buffer for 10 minutes on a wheel at 4°C. Three more washes were performed with 1ml NT2 Buffer supplemented with 0.1% Urea + 50mM NaCl (10 minutes, 4°C on a wheel). Beads were washed one more time in 500μl of NT2 buffer, 50ul were collected for WB analysis and the remaining supernatant was discarded. All the washes were performed on a magnet. RNA was extracted by adding TRIzol to the beads, according to the manufacturer’s protocol. The RNA pellets were resuspended in 15μl DEPC water and cDNAs were synthesized using the RevertAid First Strand cDNA Synthesis Kit (Thermo Fischer Scientific).

### Annexin-V Assay

Cells were seeded and treated with 10μM Nutlin for 24 or 48 hours. The FITC Annexin-V Apoptosis Detection kit I (BD Pharmingen) was used for the staining following the manufacturer’s protocol. In order to have more information about cells viability, we recovered and analyzed also cells that were in suspension after the treatments. Collected cells were then washed twice with PBS, and counted. 150000 cells were resuspended in Annexin-V binding buffer and labeled with Annexin-V and PI for 15 minutes in the dark. Finally, cells were analyzed by flow cytometry collecting 10.000 events using a BD CantoA instrument. Experiments were performed in triplicate.

### Survival curves and quantification of cell death markers using high-content imaging

U2OS cells (shNT and shDHX30 clones) were seeded in a 96-well plate format (5000 cells per well, in triplicate). Cells were imaged using digital phase contrast by an Operetta high-content imaging system (Perkin Elmer) 24 hours after seeding, just before treatment with 10μM Nutlin or DMSO control. Plates were then imaged again on the same fields at different time points. Images were processed using Harmony® High Content Imaging and Analysis Software (Perkin Elmer) to obtain survival curves. To quantify more directly the number of dead cells as a function of gene depletion (shNT vs shDHX30), treatment (DMSO vs Nutlin), and time, cells were seeded also in a 96-well plate format (5000 cells per well, in quadruplicate), stained with Propidium Iodide, and Hoechst and images acquired by Operetta were processed using Harmony® High Content Imaging and Analysis Software to quantify the numbers of total objects, the mean cell size, nuclei numbers and proportion of PI-positive cells.

