## Supplemental Figures and List of Supplemental Tables for "p53-induced apoptosis is specified by a translation program regulated by PCBP2 and DHX30"

Rizzotto, Zaccara *et al*

**Supplementary Figure Legends, list of Supplementary Tables, and Supplementary Figures**

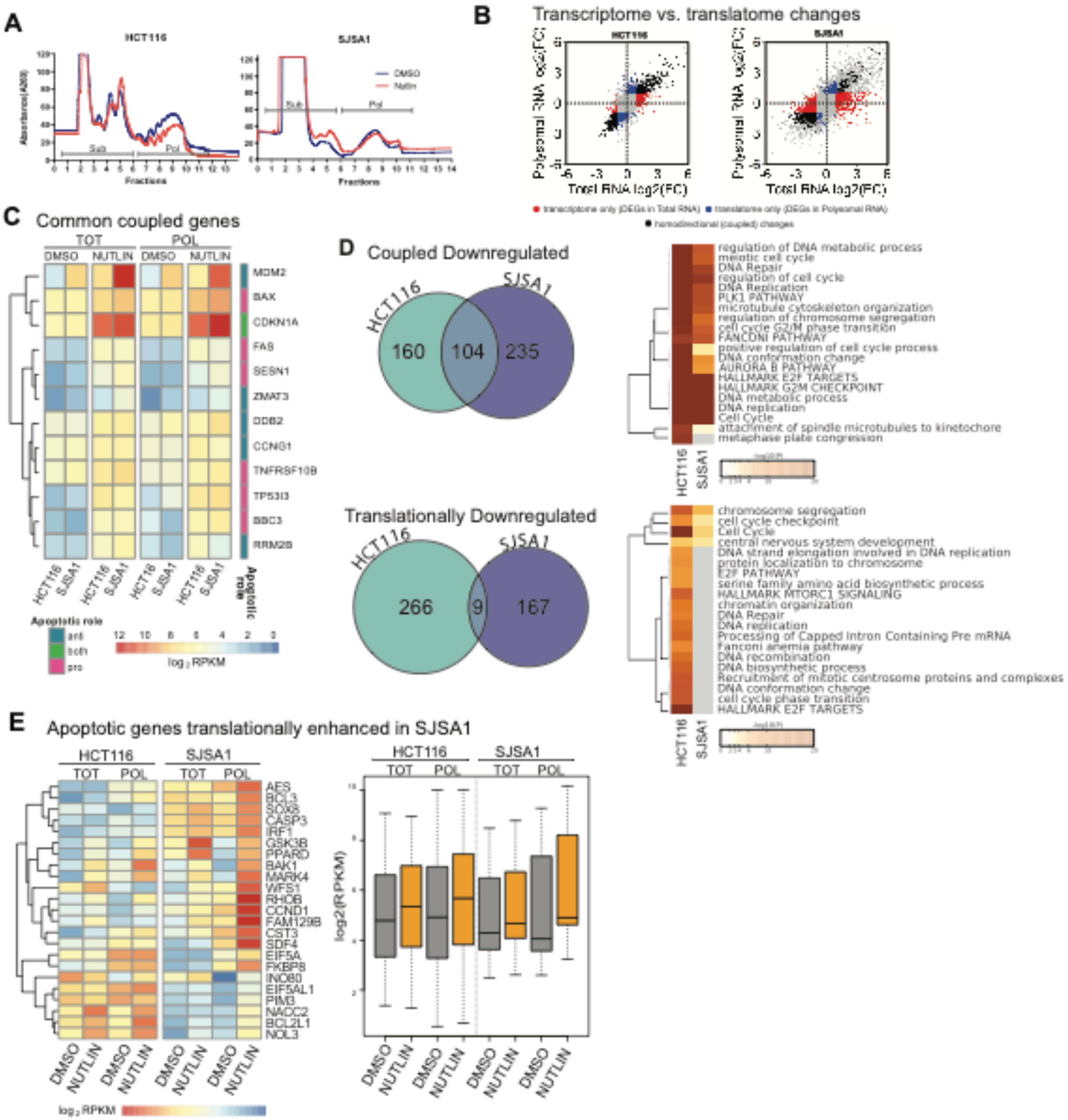


**Figure S1. Polysome profiling of HCT116 and SJSA1 cells.**

**A)** A representative image of the sucrose gradient fractionation of cytoplasmic lysates obtained by measuring the absorbance at 260nm is shown. For each cell line, a profile obtained from untreated and Nutlin treated (10µM for 12 hours) cells are shown. As depicted also in Figure 1A, fractions corresponding to two or more polysomes were collected after fractionation to extract the polysomal RNA. Likewise, the fractions corresponding to free RNA and up to the 80S monosomes, considered to be not translating, were combined to obtain the sub-polysomal RNA fraction. Nutlin treatment led to a global reduction in translation potential, as apparent from a lower amount of polysomes compared to the 80S peak, particularly evident for HCT116 cells. **B)** Scatter-plots of $log_2$ fold changes for both total and polysomal RNAs. Differentially Expressed Genes (DEGs) in each category (transcriptome only, translatome only and homo-directional changes) are classified according to $log_2$ fold change >1 and < -1 and corrected p-value <0.1 for induced and repressed genes, and colored in red, blue or black, respectively. Genes without significant changes in expression after Nutlin treatment are shown in gray. **C)** Relative expression level (expressed as read per kilobase million, $log_2$ RPKM) of a panel of well-established direct p53 targets that are common coupled DEGs according to the RNA-seq results. The attributed gene function with respect to the modulation of apoptosis is indicated. **D)** Venn diagrams of repressed DEGs in both total and polysomal RNA (Coupled Downregulated, top panel) and of DEGs that are repressed only in polysomal RNA (Translationally Downregulated, lower panel). According to GO and pathway analysis performed by Metascape (heatmaps on the right), genes related to the G2/M checkpoint and DNA damage/ repair/ replication pathways (ATM, BRCA1) are enriched among coupled as well translationally DEGs particularly in HCT116, consistent with the predominant cell cycle arrest response induced by Nutlin in these cells. **E)** Relative expression level, expressed as read per kilobase million ($log_2$ RPKM), of apoptotic genes that are translationally enhanced uniquely in SJSA1 cells responding to Nutlin treatment. RPKM values in DMSO (DMSO) and Nutlin, in total (TOT) and polysomal fractions (POL) are presented separately. A boxplot representation of the same results is included on the right to support the observed translational enhancement of apoptotic genes.

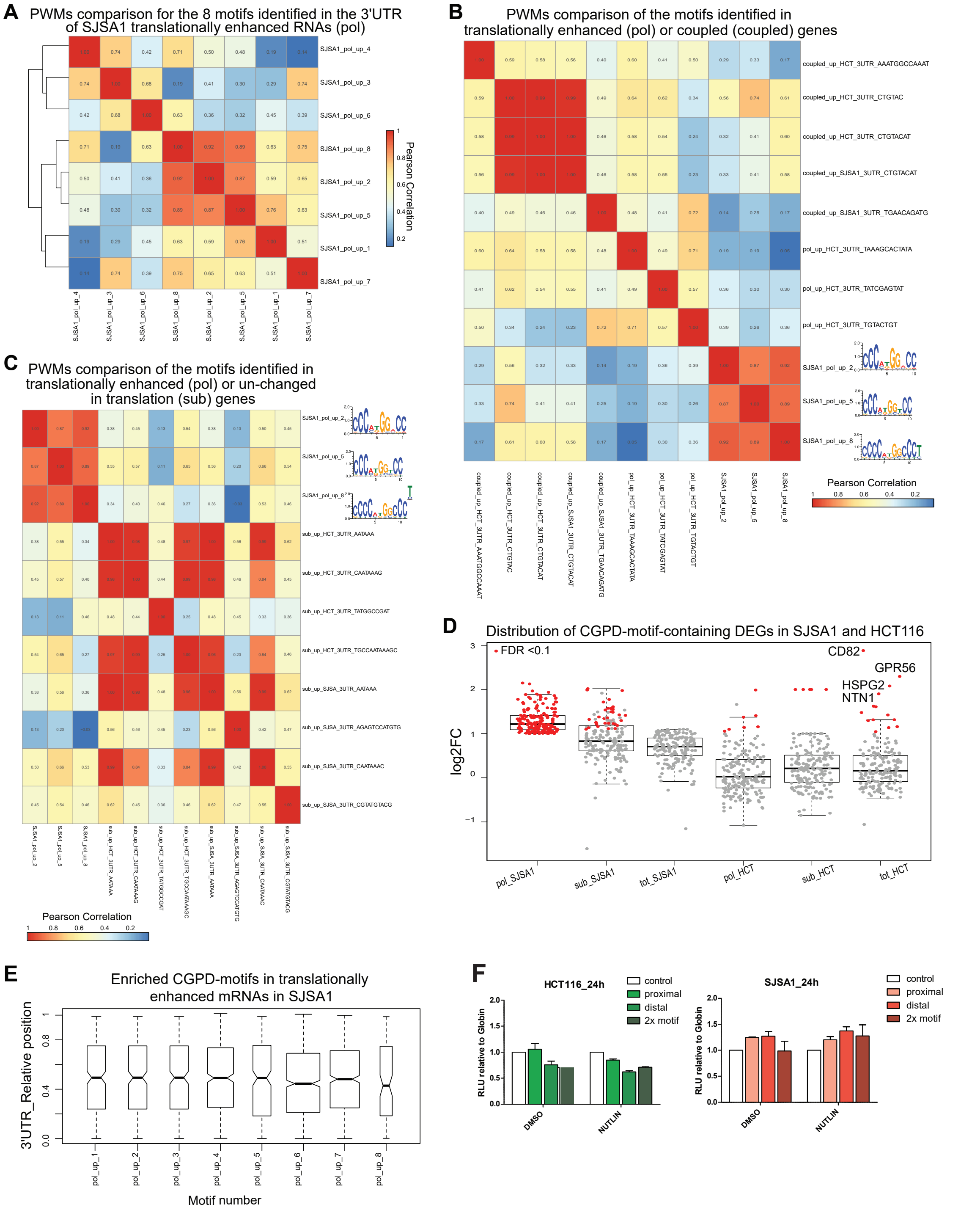


**Figure S2. Identification of a consensus motif among SJSA1 translationally enhanced mRNAs.**

All the lists of differentially expressed genes (DEGs) obtained from the RNA-sequencing of HCT116 and SJSA1 cells were investigated for the presence of over-represented cis-elements in their 3'UTRs using Weeder (see Methods for details). Motifs consensus identified by Weeder are presented in Table S4A. The level of correlation among the position weight matrices (PWMs) for the most enriched motifs for each indicated gene category is presented. **A)** Eight sequence motifs were identified as highly enriched only in SJSA1 translationally enhanced DEGs. Pearson correlation analysis indicates that three of those motifs (_2, _5, _8) are highly similar, likely representing different parts of a single extended sequence that contains a poly r(C) stretch. **B)** The comparison between enriched motifs found among coupled and translationally enhanced DEGs. The SJSA1 motifs _2, _5, _8 are exclusive for SJSA1 translated mRNAs. A CTGTAC motif is instead enriched only in coupled HCT116 mRNAs. **C)** The comparison between enriched motifs identified among translationally enhanced or un-changed in translation DEGs confirms that the SJSA1 motifs _2, _5, _8 are exclusive for SJSA1 translationally enhanced mRNAs. Common motifs (featuring A-stretches) were found between untranslated DEGs from HCT116 and SJSA1. **D)** Box plot presenting the treatment-dependent $\log_2$ fold change of the SJSA1 translated, upregulated mRNAs presenting the CGPD-motif across the other sequenced RNA fractions and in comparison, with the results from HCT116. The distribution of fold change and the absence, or the very few mRNAs that are differentially expressed (red dots) in all DEG lists besides the SJSA1 translationally regulated (pol) confirms the finding from the Pearson correlation that the CGPD-motif is enriched exclusively among SJSA1 translationally enhanced mRNAs. The list of genes that carry the motif and their $\log_2$FC in the different gene categories is presented in Table S4B. **E**) Absence of a positional bias along the 3'UTR sequence was observed for each of the SJSA1 over-represented motifs (the motifs are numbered as presented in Figure SA2). **F)** The relative light units of the reporter assay with cloned CGPD-motif within the $\beta$-globin 3'UTR from which Figure 2E is derived, are presented separately for DMSO and Nutlin treatment. A trend for constitutive down-modulation of reporter activity is visible in HCT116 cells, consistent with the hypothesis that in these cells a repressive factor exists that interacts with the CGPD-motif. The opposite trend is visible in SJSA1 cells.

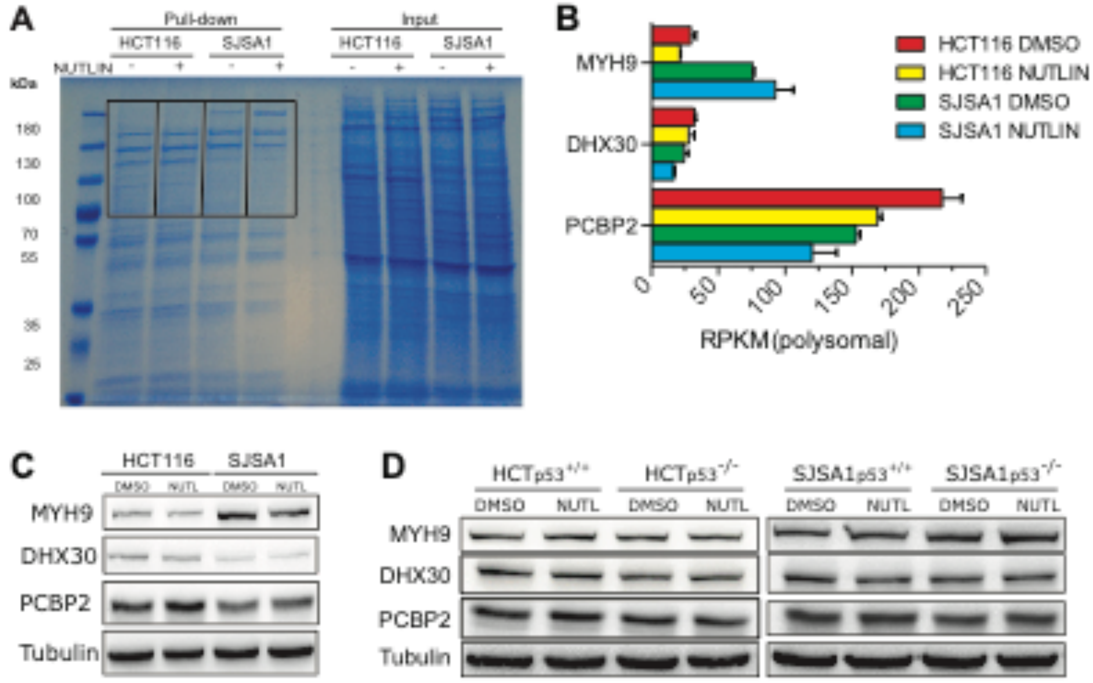


**Figure S3. Relative expression of selected proteins identified by pull-down and mass spectrometry experiments and RIP data related to Figure 4.**

**A)** SDS-PAGE of proteins extract of HCT116 or SJSA1 cells after pull-down with an RNA probe corresponding to the identified 3'UTR consensus motif linked to magnetic beads. Proteins were visualized by colloidal Coomassie blue. The visual comparison of bands from the two cellular sources of total protein and treatment was used to select a region of the gel corresponding to the ~100-180 kDa range for quantitative mass spectrometry. **B)** Relative polysomal mRNA levels (RPKM) for MYH9, DHX30, and PCBP2 according to the HCT116 and SJSA1 RNA-seq data. **C)** Relative expression of MYH9, DHX30, and PCBP2 proteins in control or Nutlin-treated HCT116 and SJSA1 cells. **D)** Comparison of MYH9, DHX30 and PCBP2 protein levels in p53 wild-type and p53 knock-out derivative clones of HCT116 and SJSA1 cells. In all western blots, Tubulin was used as a loading control.

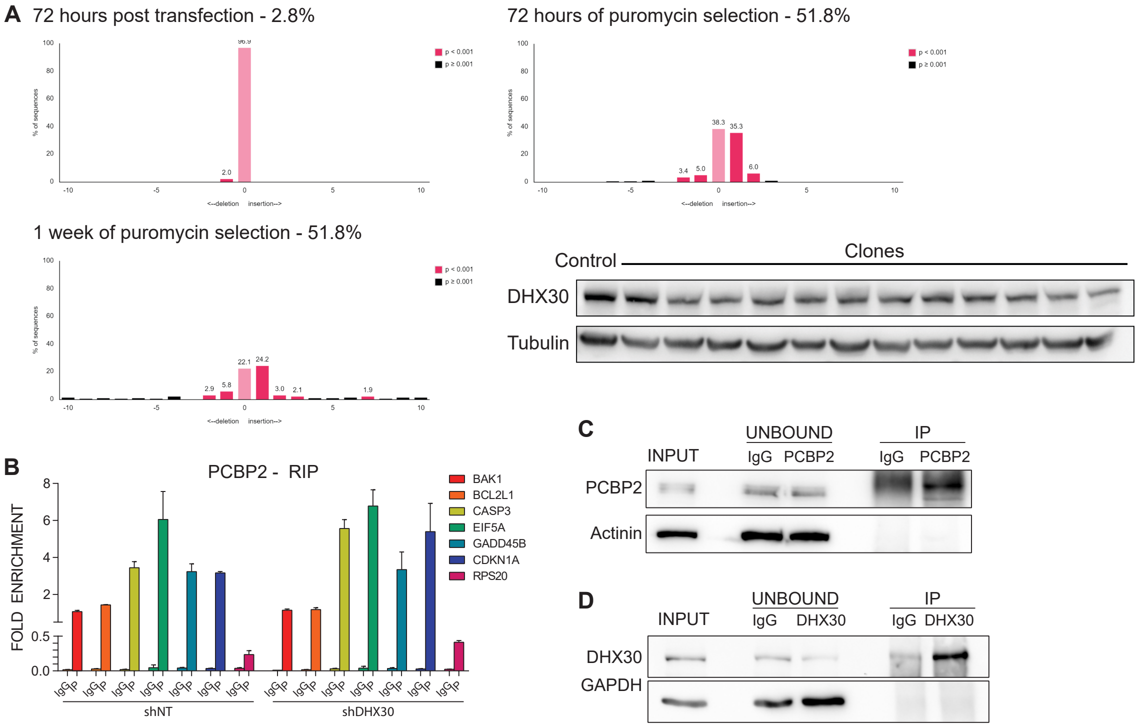


**Figure S4. CRISPR/Cas9 DHX30 knock-out experiments and RIP assays complementing Figure 4.**

**A)** Summary of indel generation efficiency in the DHX30 gene obtained by Cas9/sgRNA plasmid delivery at different time points after transfection and puromycin selection. The region surrounding the DHX30 sgRNA target site were amplified, Sanger sequenced and analyzed by TIDE web tool. Relative DHX30 expression in twelve single-clone HCT116 puromycin resistant isolated selected after transfection with the Cas9 + DHX30 guide expression plasmid. No complete knock-out clones were obtained from a total of more than fifty clones checked. **B)** The RIP protocol was performed using the PCBP2 targeting antibody starting from lysates of HCT116 shNT or shDHX30. The enrichment in immunoprecipitates are presented as % of input for the indicated CGPD-motif transcripts and control. Error bars plot the standard deviation of three technical replicates. **C)**, **D)** The efficiency of immunoprecipitation of PCBP2 and DHX30 antibody was verified by western blot, comparing unbound fractions in immunoprecipitates performed with specific antibodies or the IgG controls.

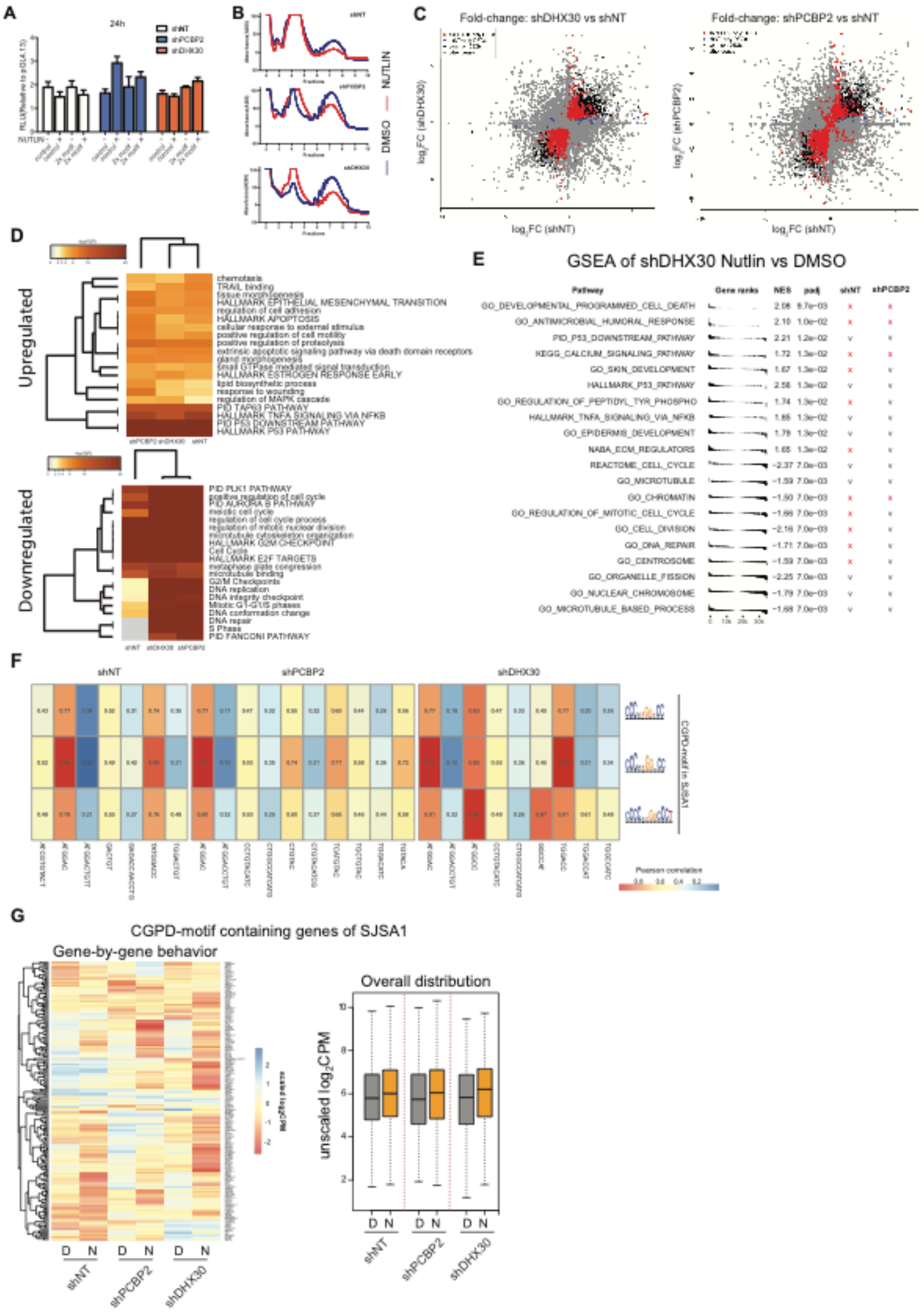


**Figure S5. Gene reporter assays, polysome profiling, RNA-sequencing and Gene Ontology analysis of HCT116 cell lines stably depleted for PCBP2, DHX30, or control.**

**A)** The relative light units obtained with the reporter construct containing two copies of CGPD-motif cloned within the β-globin 3'UTR (see Figure 5A), are presented separately for DMSO and Nutlin treatment, relative to the empty control vector. **B)** Representative image of sucrose gradient fractionation of cytoplasmic lysates of HCT116 shNT, shPCBP2 and shDHX30 cells treated or not with 10 μM Nutlin -see also Figure S1A- **C)** Scatter-plots of $\log_2$ fold changes for polysomal RNA. Left panel: comparison between shNT and shDHX30; right panel: comparison between shNT and shPCBP2. Highlighted in color are Differentially Expressed Genes (DEGs) after Nutlin treatment, defined based on the adjusted p-value threshold of 0.05. Red dots: DEGs unique for shDHX30 or shPCBP2; blue dots: DEGs unique for shNT; black dots: common DEGs to either shNT and shDHX30 or shNT and shPCBP2. Genes that are not differentially expressed are colored in gray (labeled as other genes). **D)** Metascape heatmaps summarizing GO analysis for polysomal Upregulated DEGs -top panel- or Downregulated DEGs -lower panel- from the RNA-seq data of HCT116 shNT, shPCBP2 or shDHX30 cells. **E**) Top enriched gene sets (both positive and negative Normalized Enrichment Score -NES-) for the shDHX30 Nutlin versus DMSO condition as per a GSEA. NES and adjusted p-values are shown. Enrichment status of each gene set in shNT and shPCBP2 Nutlin versus DMSO conditions are also shown (x = not enriched, v = enriched; adjusted p-value threshold = 0.05). **F)** The motifs enriched in 3'UTRs of polysome Upregulated DEGs in HCT116 shNT, shPCBP2 and shDHX30 (see Table S13) were correlated with the CGPD-motif identified in SJSA1 translated mRNAs (Figure 2A). Pearson correlation values between position weigh matrices (PWM) of enriched motifs are presented as a heatmap. **G)** Relative expression level expressed as read per kilobase million ($\log_2$ RPKM) of the 193 CGPD-motif genes previously identified in SJSA1 cells from the Weeder analysis of the Translationally Upregulated DEGs (see Figure 2A and Table S4B). Values in DMSO (D) and Nutlin (N) of polysomal fractions are presented separately. The overall distribution of the read counts is also presented as a box plot (panel on the right).

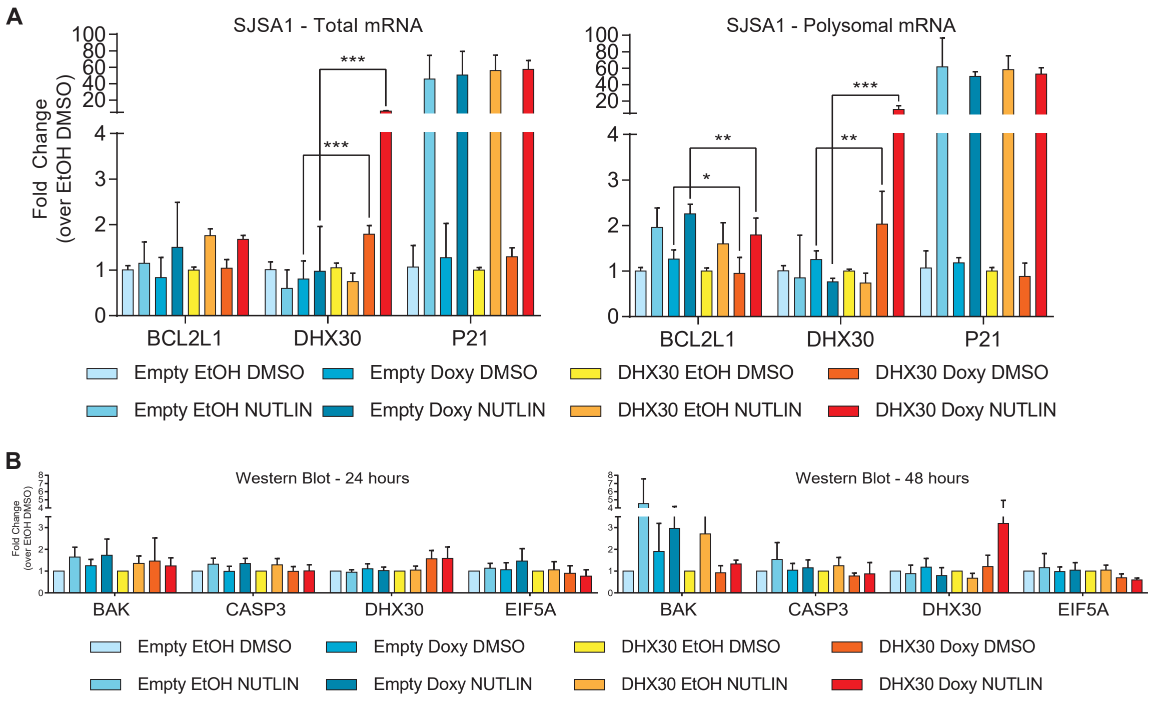


**Figure S6. Relative CGPD-mRNA and protein expression in SJSA1-DHX30-TetON cells.**

**A)** Total (left panel) or polysomal (right panel) mRNAs from SJSA1-Empty or SJSA1-DHX30 cells were used to measure the relative expression of BCL2L1, p21, and DHX30. Bars plot the average fold change relative to the EtOH treated DMSO controls. Cells were treated as described in Figure 6G. The standard deviations of at least two biological replicates are presented. (*, $p<0.05$; **, $p<0.01$; *** $p<0.001$; Student's t-test).

**B)** Bar graph presenting the relative expression of BAK, BCL2L1, CASP3, DHX30, EIF5A, and p21 proteins measured by western blot using protein extracts from SJSA1-Empty or SJSA1-DHX30 cells, exposed to 2.5µg/ml of doxycycline (or EtOH) for 24 hours prior to the treatment with Nutlin (or DMSO) for an additional 24 (left panel) or 48 (right panel) hours. Tubulin was used as loading control.

### LIST of TABLES

**Table S1**. Differentially expressed genes in HCT116 cells treated with Nutlin. Coupled, Translated, Upregulated and Downregulated lists are presented on separate sheets.

**Table S2**. Differentially expressed genes in SJSA1 cells. The file is organized as Table S1.

**Table S3.** Results of the Metascape analysis presented in Figure 1 and Figure S1.

**Table S4A**. Enriched motifs identified by Weeder in UTRs of SJSA1 DEG lists.

**Table S4B.** List of genes harboring the CGPD-motif and their relative expression levels in SJSA1 and HCT116. The expression of 182 of the 193 CGPD-motif genes is detectable also in HCT116 cells.

**Table S5.** List of RBPs identified by Mass Spectrometry as CGPD-motif interactors.

**Table S6-S8.** Differentially expressed genes in HCT116 shNT (S6), shPCBP2 (S7), shDHX30 (S8) cells filtered at adjusted p-value threshold of 0.05. Tables also contain the lists of unique apoptotic genes that are translationally regulated (see Figure 5C).

**Table S9**: Results of the Metascape analysis presented in Figure S5.

**Table S10-S12:** Enriched gene sets as per the GSEA of shNT (S10), shPCBP2 (S11), shDHX30 (S12), Nutlin vs DMSO condition filtered at adjusted p-value threshold of 0.05.

**Table S13**. Enriched motifs identified by Weeder for HCT116 shNT, shPCBP2, and shDHX30 RNA-seq data.

**Table S14:** List of primers.

**Table S15**: Quantification of western blot bands. The worksheet name indicates the Figure panels that were quantified using UVITEC Alliance (UVITEC Cambridge). Average relative protein amounts and standard deviation of three replicates are presented as a table and also, as summary graphs. Tubulin was quantified as a loading control.
